# Disposition and Metabolomic Effects of 2,2’,5,5’-Tetrachlorobiphenyl in Female Rats Following Intraperitoneal Exposure

**DOI:** 10.1101/2023.06.19.544952

**Authors:** Amanda Bullert, Xueshu Li, Chunyun Zhang, Kendra Lee, Casey F. Pulliam, Brianna S. Cagle, Jonathan A. Doorn, Aloysius J. Klingelhutz, Larry W. Robertson, Hans-Joachim Lehmler

## Abstract

The disposition and toxicity of lower chlorinated PCBs (LC-PCBs) with less than five chlorine substituents have received little attention. This study characterizes the distribution and metabolomic effects of PCB 52, an LC-PCB found in indoor and outdoor air, three weeks after intraperitoneal exposure of female Sprague Dawley rats to 0, 1, 10, or 100 mg/kg BW. PCB 52 exposure did not affect overall body weight. Gas chromatography-tandem mass spectrometry (GC-MS/MS) analysis identified PCB 52 in all tissues investigated. Hydroxylated, sulfated, and methylated PCB metabolites, identified using GC-MS/MS and nontarget liquid chromatography-high resolution mass spectrometry (Nt-LCMS), were primarily found in the serum and liver of rats exposed to 100 mg/kg BW. Metabolomic analysis revealed minor effects on L-cysteine, glycine, cytosine, sphingosine, thymine, linoleic acid, orotic acid, L-histidine, and erythrose serum levels. Thus, the metabolism of PCB 52 and its effects on the metabolome must be considered in toxicity studies.

**Highlights:**

- PCB 52 was present in adipose, brain, liver, and serum 3 weeks after PCB exposure
- Liver and serum contained hydroxylated, sulfated, and methylated PCB 52 metabolites
- Metabolomics analysis revealed minor changes in endogenous serum metabolites
- Levels of dopamine and its metabolites in the brain were not affected by PCB 52

## 1. Introduction

Polychlorinated biphenyls (PCBs), a class of 209 industrial chemicals, were manufactured and sold as complex mixtures of over 150 individual PCB congeners. Because of their physicochemical properties, they were used in various commercial applications, including insulating fluids in transformers, building materials, adhesives, and resins (ATSDR, 2000; Faroon and Ruiz, 2016; Markowitz and Rosner, 2018). The industrial production of PCBs has been banned worldwide because of their environmental persistence, long-range transport in the environment, the potential to bioaccumulate and biomagnify in aquatic and terrestrial food chains, and environmental and human health concerns (Stockholm-Convention, 2019). PCBs have been released into the environment and can be found, for example, in soils, sediments, water sources, wildlife, and air (Hu et al., 2011; Marek et al., 2017). In addition, PCBs continue to be inadvertently produced by various industrial processes and can be found, for example, in certain paints and adhesives used in kitchen cabinets (Anezaki and Nakano, 2013; Herkert et al., 2018; Hu and Hornbuckle, 2010).

PCBs and their metabolites are present in human tissues because of the widespread environmental contamination with PCBs (Ampleman et al., 2015; Li et al., 2022; Sethi et al., 2019). Exposure of the general population to PCBs primarily occurs via contaminated foods like fish, meats, and dairy (Saktrakulkla et al., 2020; Schecter et al., 2010). In addition, growing evidence demonstrates that inhalation of indoor or outdoor air represents a current route of human PCB exposure (Ampleman et al., 2015; Harrad et al., 2006; Kaifie et al., 2019; Meyer et al., 2013). Of particular concern is the presence of PCBs in the indoor air of schools (Egsmose et al., 2016; Marek et al., 2017). Therefore, exposures to PCBs represent a current human health concern (Agus et al., 2022; Hammel et al., 2023). PCB exposure has been linked to cancer and non-cancer endpoints (Carlson et al., 2022; 2016). Importantly, laboratory and population-based studies implicating higher chlorinated PCBs (i.e., congeners with > 4 chlorine substituents) and their metabolites in neurotoxic outcomes across the lifespan (Bullert et al., 2021; Gaum et al., 2014; Klocke and Lein, 2020; Pessah et al., 2010). Lower chlorinated PCBs have not been studied as extensively for their neurotoxic effects as compared to their association with cancer and other outcomes (Deen et al., 2023; Hammel et al., 2023; Kofoed et al., 2021).

PCB 52, a tetrachlorinated biphenyl, is a major constituent of technical PCB mixtures (Frame, 1997; Frame et al., 1996) and has been widely detected in air and human samples (Grimm et al., 2015b; Hammel et al., 2023). This tetrachlorinated PCB congener is metabolized via an arene oxide intermediate and by direct insertion of oxygen into a C-H bond to 3-OH-PCB 52, 4-OH-PCB 52, and 4,5-diOH-PCB 52 by rat liver microsomes (Borlakoglu et al., 1991; Borlakoglu and Wilkins, 1993; Forgue et al., 1979). CYP2A6 is the cytochrome P450 (CYP) isoform that forms 4-OH-PCB 52 in humans (Shimada et al., 2016). The CYPs involved in the metabolism of PCB 52 in rodents are unknown; however, PCB 52 is likely oxidized by rats CYP2B1, similar to PCB 95 and PCB 136 (Lu et al., 2013; Lu and Wong, 2011). OH-PCB 52 metabolites can undergo further oxidation, glucuronidation, and sulfation, resulting in complex metabolite mixtures (Grimm et al., 2015b). Although monohydroxylated PCB 52 metabolites have not been detected in human samples, the corresponding PCB 52 sulfate metabolites have been recently detected in human populations (Zhang et al., 2022b).

PCB metabolites can exhibit toxicity, including endocrine disruption and carcinogenic and neurologic effects. For example, OH-PCBs interact with estrogen receptors as either agonists or antagonists, with some OH-PCBs being more effective agonists than estradiol based on their interaction with the estrogen receptors (Grimm et al., 2015b; Tehrani and Van Aken, 2014). 4-OH-PCB 52 was found to have a low affinity for the thyroid hormone receptor and display low estrogenic activity *in vitro* (Kitamura et al., 2005). PCBs and OH-PCBs can be further metabolized to potentially toxic PCB sulfates by sulfotransferase enzymes (SULT) (Grimm et al., 2015b). 4-OH-PCB 52 and the corresponding sulfate metabolite are inhibitors of SULT2A1, while the PCB 52 sulfate is an inhibitor of SULT1E1 (Parker et al., 2018). In addition, OH-PCB 52 and its PCB 52 sulfate displayed toxicity in the low micromolar range in two neural cell lines, the N27 and SH-SY5Y cell lines (Rodriguez et al., 2018). PCB 52 can also be metabolized to redox reactive quinone metabolites, as demonstrated by the presence of quinonoid-derived protein adducts in the liver and brain of PCB 52 exposed Sprague-Dawley rats (Lin et al., 2000). PCB quinones and other PCB metabolites can lead, for example, to gene mutations, inhibition of nuclear protein topoisomerase, and a reduction in telomerase activity (Grimm et al., 2015b; McLean et al., 1996; Song et al., 2008).

This exploratory study characterizes the disposition of PCB 52 and its metabolites in female Sprague Dawley rats exposed to different doses of PCB 52 using both targeted and nontargeted approaches and assesses changes in endogenous metabolites implicated in PCB neurotoxicity. Our results reveal minor the transformation of PCB 52 to a complex metabolite mixture in rats.

## 2. Materials and Methods

### 2.1. Chemicals

The sources of the test compound, PCB 52, and the analytical standards are described in the Supplementary Material. Unique chemical identifiers are summarized in **Table S1**. Pesticide grade solvents, including hexane and dichloromethane, and mass spectrometry grade solvent, such as acetonitrile, were purchased from Fisher Scientific.

### 2.2. Animal exposure

All experiments were conducted with the approval of the Institutional Animal Care and Use Committee of the University of Iowa. Female Sprague Dawley (SD) rats, weighing 75–100 g at 4–5 weeks old, were supplied by Harlan Sprague-Dawley/Envigo (Indianapolis, Indiana, USA). The animals were provided a standard rodent chow and water *ad libitum*. After a week of acclimation, rats were administered a single intraperitoneal (IP) injection of corn oil (vehicle control) or PCB 52 in corn oil at PCB 52 doses of 1 mg/kg, 10 mg/kg, or 100 mg/kg BW (body weight). While these doses are higher than the Tolerable Daily Intake (TDI) of PCBs for humans of 1.6 µg per day for an 80 kg individual (Faroon, 2003), the use of high doses and short study periods is a well-established approach for generating data useful for risk assessment purposes (Doull, 2003). Three weeks after PCB 52 administration, all rats were euthanized using carbon-dioxide asphyxiation followed by thoracotomy. After euthanasia, whole blood was obtained from the heart and allowed to rest at room temperature for at least 30 min, followed by centrifuge at 2,500 g for 10 min at 4°C to prepare serum. The serum was then aliquoted into glass vials. The livers, spleen, and thymus were excised *en bloc*, flash-frozen in liquid nitrogen. All samples were stored at −80°C until further analysis.

### 2.3. Targeted GC-MS/MS analysis of PCB 52 and metabolites

#### 2.3.1. Extraction of PCB 52 and metabolites from tissues

PCB 52 and its hydroxylated metabolites were extracted from the adipose (109 ± 13 mg range: 95 – 139 mg), brain (648 ± 77 mg range: 460 – 760 mg), and liver (509 ± 4 mg range: 503 –515 mg) using the “Jensen” extraction method (Jensen et al., 2003). Briefly, tissues were homogenized in 3 mL of isopropanol with 1 mL of diethyl ether, as described (Kania-Korwel et al., 2008b; Wu et al., 2015). Recovery standards (PCB 77, 20 ng in isooctane; 4’-OH-PCB 159, 20 ng in methanol) were added to each sample. The samples were capped, inverted for 5 min, and centrifuged at 1,690 g for 5 min. Next, the supernatant was transferred to a new glass tube containing 5 mL of 0.1 M phosphoric acid in 0.9% NaCl solution. The tissue pellets were resuspended with 1 mL of isopropanol and 2.5 mL hexane-diethyl ether (9:1, v/v), followed by vortexing. Samples were then inverted and centrifuged as described above, and the supernatant was combined with the previous extract. Extracts were inverted and centrifuged as described above to facilitate phase separation. The organic phase was transferred to a new vial while the aqueous phase was re-extracted with 3 mL of the hexane-diethyl ether solution (9:1, v/v). The combined organic phases were concentrated under a slow nitrogen stream.

Five drops of methanol and 0.5 mL of diazomethane (about 5 mmol of diazomethane) in diethyl ether were added to the tissue extracts (Kania-Korwel et al., 2008b). The samples were stored at 4° C for at least 3 h to methylate hydroxylated PCB 52 metabolites. After derivatization, excess diazomethane was evaporated under a gentle stream of nitrogen, and samples were loaded onto glass solid phase extraction (SPE) cartridges containing 0.2 g of activated silica gel (bottom) and 2 g of acidified silica gel (silica gel: H_2_SO_4_, 2 :1, w/w; top). Analytes were eluted with 10 mL of dichloromethane, and the extracts were concentrated to near dryness under a gentle nitrogen stream. The tissue extracts were treated with 4 mL of concentrated sulfuric acid for further lipids removal. The organic phase was transferred to a new vial, concentrated to near dryness, and samples were spiked with the internal standard (PCB 204, 20 ng in isooctane) for gas chromatographic analysis.

#### 2.3.2. Extraction of PCB 52 and metabolites from serum for GC-MS/MS analysis

Serum (342 ± 92 mg range: 93 – 477 mg) samples were processed using a previously validated extraction protocol (Li et al., 2019). Briefly, 3 mL of aqueous 1% KCl solution was added to each serum sample, followed by the recovery standards (PCB 77, 20 ng in isooctane; 4’-OH-PCB 159, 20 ng in methanol). Subsequently, 1 mL of 6 M HCl, 5 mL of 2-propanol, and 5 mL of a hexanes-MTBE mixture (1:1 v/v) were added to each sample, vortexing between each solvent addition. The samples were inverted and centrifuged to facilitate separation between the organic and aqueous phases. The organic phases were transferred to a new glass tube, while the aqueous phase was re-extracted with 3 mL of hexanes. The combined organic phase was washed with 3 mL of aqueous 1% KCl and concentrated under a gentle stream of nitrogen. The extracts were diluted with 0.5 mL, derivatized with diazomethane as described above for tissue extracts, and concentrated with a gentle nitrogen stream. The internal standard (PCB 204, 20 ng in isooctane) was added before the gas chromatographic analysis. Quality assurance samples were analyzed in parallel with all samples, as described for the tissue samples.

#### 2.3.3. GC-MS/MS analysis of PCB 52 and its metabolites

PCB 52 and its mono-hydroxylated metabolites (as methylated derivatives) were quantified using the multiple reaction monitoring setting (MRM) on an Agilent 7890 A GC system equipped with an Agilent 7000 Triple Quad and an Agilent 7693 autosampler (GC-MS/MS) from Agilent Technologies, Inc. (Santa Clara, CA, USA). Separations were performed with an SPB-Octyl capillary column (30 m length, 25 mm inner diameter, 0.25 μm film thicknesses: Supelco, Bellefonte, Pennsylvania, USA). Samples were injected in the solvent vent injection mode with a helium (carrier gas) flow of 0.75 mL/min and nitrogen as the collision gas. The temperature program was set as follows: Initial temperature of 45 °C, hold for 2 minutes, 100 °C/min to 75 °C, hold for 5 mins, 15 °C/min to 150 °C, hold for 1 minute, 2.5 °C/min to 280 °C, and final hold of 5 minutes. Precursor and product ions of all analytical standards and the corresponding collision energy are summarized in **Table S2**.

#### 2.3.4. GC-MS/MS analysis quality assurance and quality control

All PCB 52 analyses followed validated standard operating procedures (SOPs) to extract and quantify PCB 52 and its hydroxylated metabolites. Surrogate recovery standards, PCB 77 and 4’-OH-PCB 159, were added to each sample to assess the precision and reproducibility of the PCB and OH-PCB measurements. The levels of PCB 52 and its primary metabolite, 4-OH-PCB 52, were corrected based on the recoveries of the corresponding recovery standard (**Table S3**). In parallel, the precision and recoveries of the analyses were assessed by spiking sample blanks and matched blank tissue matrices with known amounts of PCB 52 and 4-OH-PCB 52 (**Table S4**). Detection limits were determined from the method blanks or blank tissues analyzed in parallel with the samples (**Table S5**).

### 2.4. Nontarget liquid chromatography-high resolution mass spectrometric analysis (Nt-LCMS)

#### 2.4.1. Extraction of PCB 52 and metabolites from serum or tissues

Brain (213 ± 4 mg, range: 203 – 217 mg), liver ( 249 ± 2 mg, range: 245 – 254 mg), and serum samples (127 ± 57 mg, range: 51 – 282 mg) were processed using a QuEChERS method, as described with modifications (Li et al., 2022; Li et al., 2021). Briefly, brain or liver tissue was homogenized using a TissueRuptor from Qiagen (Germantown, Maryland, USA) in 2 mL of water, followed by adding the surrogate standard, F-OH-PCB 3 (50 ng in acetonitrile). Similarly, serum was added to 2 mL of water and vortexed, followed by the addition of the surrogate standards (50 ng each). After vortexing, 4 mL of acetonitrile was added. Then, a mixture of 200 mg of NaCl and 800 mg of MgSO_4_ was added, and samples were capped and shaken vigorously. Samples were then inverted for 5 minutes and centrifuged (1811 g) for 5 minutes to facilitate phase separation.

Hybrid phospholipid solid phase extraction (HybridSPE) cartridges (100 mg/3 mL, Sigma-Aldrich, St. Louis, Missouri, USA) were modified by adding 3 g of a MgSO_4_:Na_2_SO_4_ mixture (1:1, w:w) and preconditioned with 3 mL of acetonitrile. The acetonitrile layer from the QuEChERS extraction was then passed through the HybridSPE cartridge. The aqueous phase was re-extracted with 1 mL of acetonitrile, vortexed, and centrifuged for 5 minutes. The organic layer from the second extraction step was also passed through the HybridSPE cartridge, and the metabolites were eluted using 3 mL of acetonitrile. Next, the samples were evaporated to dryness at 35 °C using a Savant SpeedVac SPD210 (ThermoFisher Scientific, Waltham, Massachusetts, USA). Next, the samples were reconstituted in 300 µL of acetonitrile, vortexed, and centrifuged for 5 minutes. The supernatants were transferred to a microcentrifuge tube and spiked with perfluorooctanesulfonic acid potassium salt (50 ng). The solvent was again evaporated to dryness at 35 °C using the SpeedVac. The resulting extracts were reconstituted with 200 µL of water:acetonitrile (1:1, v/v) and vortexed. Finally, the samples were kept at −20 °C for at least 30 minutes, centrifuged repeatedly at 4 °C for 10 minutes at 16,000 g, and stored at −80 °C before Nt-LCMS analysis.

#### 2.4.2. Liquid chromatography-high resolution mass spectrometry analysis

Serum and tissue extracts were analyzed on a Q-Exactive Orbitrap Mass Spectrometer (ThermoFisher Scientific) with a Vanquish Flex ultra-high-performance liquid chromatograph (ThermoFisher Scientific) with an ACQUITY UPLC-C18 column (particle size: 1.7 µm, 2.1 x 100 mm, Waters, Milford, Massachusetts, USA) at the High-Resolution Mass Spectrometry Facility of the University of Iowa (Zhang et al., 2020). Water and acetonitrile were used as mobile phase A and B, with a flow rate of 0.3 mL/min. The pressure of the chromatographic system ranged from 4000 to 8000 psi. The UHPLC gradient program was as follows: starting at 5% B, held for 1 min, increased linearly to 95% B, held for 3 min, and returning to 5% B, with a hold for 4 min before the next injection. The injection volume was 2 µL. The Q-Exactive Orbitrap Mass Spectrometry was operated in the negative polarity mode. The spray voltage and current were 2472 V and 18.2 µA. The flow rate of sheath and auxiliary gas were 48 and 2 mL/min, respectively. The capillary and auxiliary temperature were 256 °C and 413 °C, respectively. The analyses were performed in the full scan mode with a range of 85 to 1000 m/z. The autogain control target setting was 1×106, the full scan resolution setting was 70,000 and maximum interval time (IT) was 200 ms.

#### 2.4.3. Nt-LCMS analysis quality assurance and quality control

Solvent (50% water/50% acetonitrile) and method blanks were used in the Nt-LCMS to monitor carryover. No OH-PCB metabolites were detected in these blank samples. The average recoveries of the surrogate standard (i.e., 3-F-PCB3 sulfate) were 74 ± 18% (range 45-130%, N = 47).

#### 2.4.4. Processing and visualization of Nt-LCMS data

The chromatographic peaks of suspected PCB metabolites were extracted from the acquired data (as .raw files) with Thermo Xcalibur (version 4.1, Thermo Fisher Scientific, USA). The chromatogram extraction was performed with a m/z tolerance of 5 ppm, mass precision decimals of 5, and a smoothing factor of 7. The isotopic pattern of chlorine was used as an important factor for confirming putative chlorine compounds, as described (Li et al., 2022). The chromatographic and mass spectrometric data were exported, and figures were prepared with GraphPad Prism (version 9.4.1, GraphPad Software, USA).

### 2.5. Targeted metabolomics of serum samples

#### 2.5.1. Sample preparation

Targeted metabolomic analyses were performed by the Metabolomics Core Facility of the Fraternal Order of Eagles Diabetes Research Center at the University of Iowa (Iowa City, Iowa, USA) using standard protocols (Tompkins et al., 2019). Briefly, the serum samples were extracted in ice-cold 1:1 methanol/acetonitrile containing a mixture of nine internal standards (d_4_-Citric Acid, ^13^C_5_-Glutamine, ^13^C_5_-Glutamic Acid, ^13^C_6_-Lysine, ^13^C_5_-Methionine, ^13^C_3_-Serine, d_4_-Succinic Acid, ^13^C_11_-Tryptophan, d_8_-Valine; Cambridge Isotope Laboratories, 1 μg/mL each). The ratio of extraction solvent to sample volume was 18:1. The samples were then incubated at – 20°C for 1 h, followed by a 10-minute centrifugation. Supernatants were transferred to fresh tubes. Pooled quality control (QC) samples were prepared by adding an equal volume of each sample to a fresh 1.5 mL microcentrifuge tube. Processing blanks were created by adding extraction solvent to microcentrifuge tubes. Serum samples, QC samples, and processing blanks were evaporated using a Savant SpeedVac SPD210 (ThermoFisher). The resulting dried extracts were derivatized using methyoxyamine hydrochloride (MOC) and N,O-bis(trimethylsilyl)trifluoroacetamide (TMS) (Sigma-Aldrich, St. Louis, MO, USA). The dried extracts were reconstituted in 60 μL of 11.4 mg/mL MOC in anhydrous pyridine (VWR International, Radnor, PA, USA), vortexed for 5 min, and heated for 1 hour at 60°C. Next, 40 μL TMS was added to each sample, and samples were vortexed for 1 minute before heating for 30 minutes at 60°C. The serum, QC, and blank samples were immediately analyzed using GC-Orbitrap MS.

#### 2.5.2. Targeted analysis of serum sample using GC-Orbitrap MS

Gas chromatographic separation was conducted on a ThermoFisher Scientific Trace 1300 GC with a TraceGold TG-5SilMS column (0.25 μm film thickness; 0.25 mm ID; 30 m length; ThermoFisher Scientific), as described (Tompkins et al., 2019). The injection volume of 1 μL was used for all samples. The gas chromatograph was operated in split mode with the following settings: 20:1 split ratio; split flow: 24 μL/min, purge flow: 5 mL/min, Carrier mode: Constant Flow, Carrier flow rate: 1.2 mL/min). The inlet temperature was 250°C. The oven temperature gradient was: 80°C for 3 minutes, ramped at 20°C/minute to a maximum temperature of 280°C, held for 8 minutes. The injection syringe was washed 3 times with methanol and 3 times with pyridine between samples. Metabolites were detected using a ThermoFisher Scientific Q-Exactive Plus mass spectrometer. The data were acquired from 3.90 to 21.00 minutes in EI mode (−70eV) in full scan (m/z 56.7-850) at 60K resolution. Metabolite profiling data were analyzed using Tracefinder 4.1, utilizing standard verified peaks and retention times.

#### 2.5.3. Analysis of targeted serum metabolomics data

Targeted metabolomic data were collected with a target list containing 94 metabolites. Metabolites with > 50 % missing values were removed from further analysis. The remaining missing values were imputed with the one-half lowest intensity detected for those metabolites across all samples. Metabolites detected with a relative standard deviation > 30 % in multiple injected QC samples were removed from the dataset. As a result, 81 metabolites were included for further data processing. The data were normalized by the sum of total metabolite intensity and log2 transformed. Data variability and group clustering were visualized with score plots in principal component analysis (PCA) and partial least squares-discriminant analysis (PLS-DA). Metabolites that drive the sample clustering among exposure groups were indicated with the variable importance in projection (VIP) values in the PLS-DA. Further univariate analysis was performed with a linear regression model using the *LinearModelFit* function in the R package *NormalizeMets* (De Livera et al., 2018) to generate raw p-values reflecting the correlation between metabolite intensities and dose levels. Raw p-values were adjusted for the false discovery rate with the Benjamini and Hochberg method (Benjamini and Hochberg, 1995). Analyses were performed with MetaboAnalyst 4.0 (Chong et al., 2018) or R (version 4.1.2). No pathway analysis was performed since only a limited number of metabolites (9 out of 81) were obtained with raw p-values < 0.05.

### 2.6. Dopamine and metabolites analysis

Brains were isolated and nigrostriatal tissue collected for further analysis. Tissue was homogenized in ice-cold 0.3 M perchloric acid and stored at −80°C. Prior to HPLC analysis, the homogenate was centrifuged at 10,000 g for 10 min and supernatant injected into the HPLC for neurochemistry analysis. The weight of tissue collected was used to normalize the pmol of dopamine and dopamine metabolites detected to mg of tissue. Dopamine and its metabolites, 3,4-dihydroxyphenylacetic acid (DOPAC) and 3,4-dihydroxyphenylacetaldehyde (DOPAL), were analyzed via an Agilent 1100 Series capillary HPLC system with an ESA Coulochem III coulometric electrochemical detector (Enayah et al., 2018). Separation was achieved with a Phenomenex Synergi C18 column (2 × 150 mm, 40 Å) using an isocratic mobile phase (50 mM citric acid, 1.8 mM sodium heptane sulfonate, 0.2% trifluoroacetic acid, 2% acetonitrile, pH = 3.0) at 200 μL/min. For electrochemical detection of catechol-containing compounds, the guard cell, electrode 1, and electrode 2 were set to +350, −150, and +200 mV, respectively.

### 2.7. Statistical analysis

Data are presented as means ± standard deviation. The differences among means were compared using one-way or two-way analysis of variance (ANOVA) with Tukey’s post hoc method testing in GraphPad Prism (version 9.0.0.), as appropriate. P-values less than 0.05 were considered statistically significant. The original data underlying this study are openly available through the Iowa Research Online repository at DOI: 10.25820/data.006225.

## 3. Results and Discussion

### 3.1. General toxicity

Exposure to PCB 52 had no significant impact on the overall body weight of the adolescent rats (**Fig. 1a**). In contrast, other studies have shown that PCB exposure increases mammalian body weights during critical growth phases, such as adolescence (Birnbaum, 1994; Corey et al., 1996). This increase is due to altered hormone homeostasis, inflammation of organs, and increasing organ weights affecting overall body weight following PCB exposure. PCB 52 exposure did not affect spleen and thymus weights (**Figs. S1** and **S2**), while the bodyweight-adjusted livers of animals exposed to 10 mg/kg BW of PCB 52 were significantly smaller than the vehicle group (**Fig. 1b**). In an earlier study, bodyweight-adjusted liver weights from male rats exposed to a single IP dose (150 mg/kg BW) of PCB 52 where not significantly different than their control counterparts (**Table S6)** (Parkinson et al., 1983). In contrast, exposure to many PCB mixtures or higher chlorinated PCBs typically results in an enlarged liver (ATSDR, 2000; Parkinson et al., 1983).

**Fig. 1.**
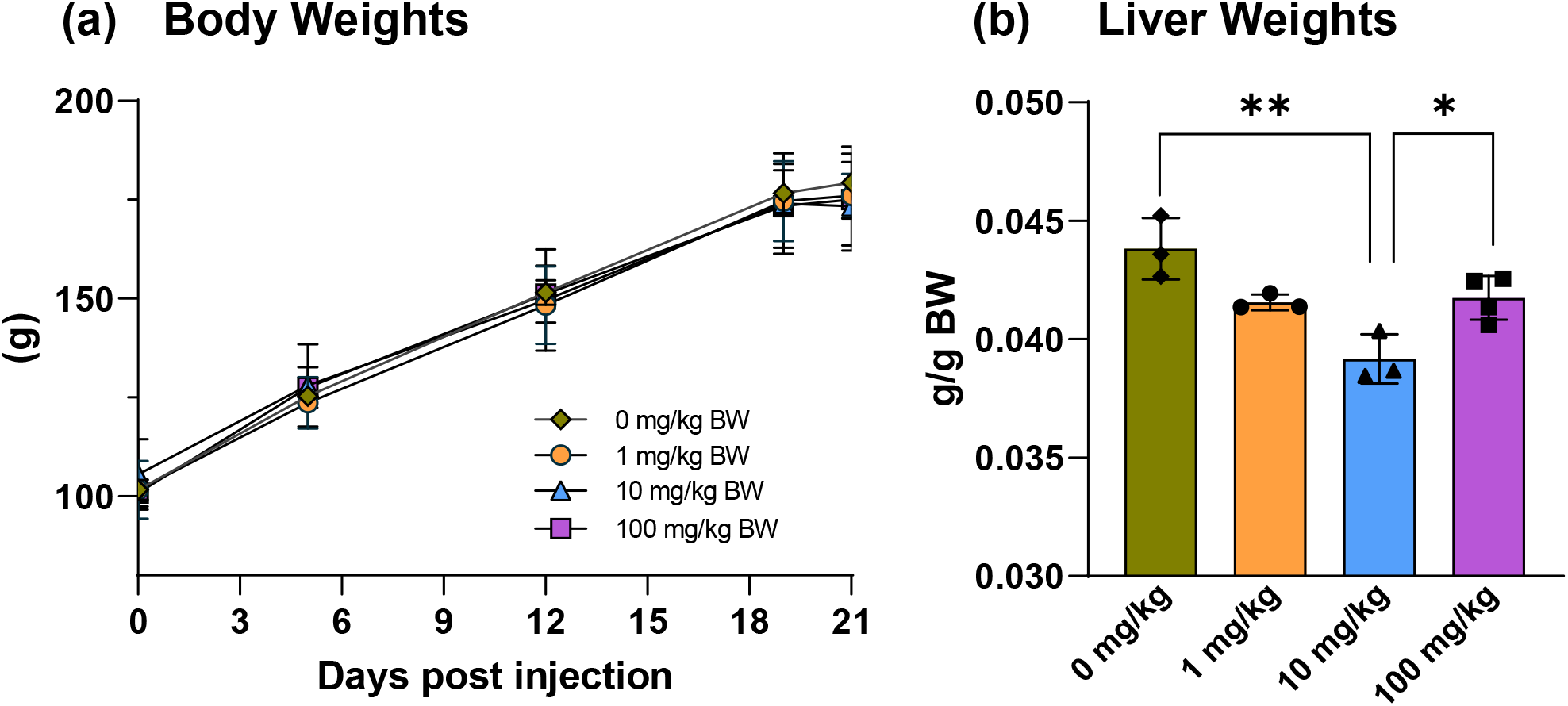
PCB 52 exposure did not affect body weights but altered relative liver weights following intraperitoneal (IP) exposure of female rats to different doses of PCB 52. (A) Growth curve of body weights throughout the 3-week study, measured at 5 different time points post IP injection of PCB 52. (B) Body weight adjusted liver weights. A significant decrease in liver weights was measured 3 weeks post IP injection of PCB 52 for rats exposed to 10 mg/kg BW. Group means were compared using one-way ANOVA followed by Tukey’s pairwise comparisons. A p-value < 0.05 was considered statistically significant.

### 3.2. PCB tissue distribution

PCB 52 is a potentially neurotoxic PCB congener that, based on congener-specific PCB analyses, is frequently detected in air and human serum form the United States (summarized in (Grimm et al., 2015b). PCB 52 was present in all tissues investigated, including the brain, three weeks following the IP injection (**Fig. 2, Table S7**). Except for the high dose group, PCB 52 levels followed the rank order adipose > liver ∼ brain ∼ blood across all three exposure groups. In the high-dose group, the mean PCB 52 levels in the liver were higher than in the brain and blood. Disposition studies with PCB 95, a PCB congener structurally related to PCB 52, show a similar rank order in mice, with adipose ≫ liver > brain > blood (Kania-Korwel et al., 2015; Kania-Korwel et al., 2012). In the present study, PCB 52 levels were 50 to 100 times higher in the adipose tissue than in all other tissues investigated. The high PCB 52 levels in adipose tissue, the primary storage site of PCBs (Kania-Korwel et al., 2008a), are due to its high triglyceride content compared to the other tissues (McCann et al., 2021; Seelbach et al., 2010; Tan et al., 2004). PCB 52 was also detected in the liver, the primary site of oxidative PCB metabolism, in part because the blood vessels of the peritoneum drain directly into the liver, contributing to a sustained release of PCB 52 into the liver (Al Shoyaib et al., 2019).

**Fig. 2.**
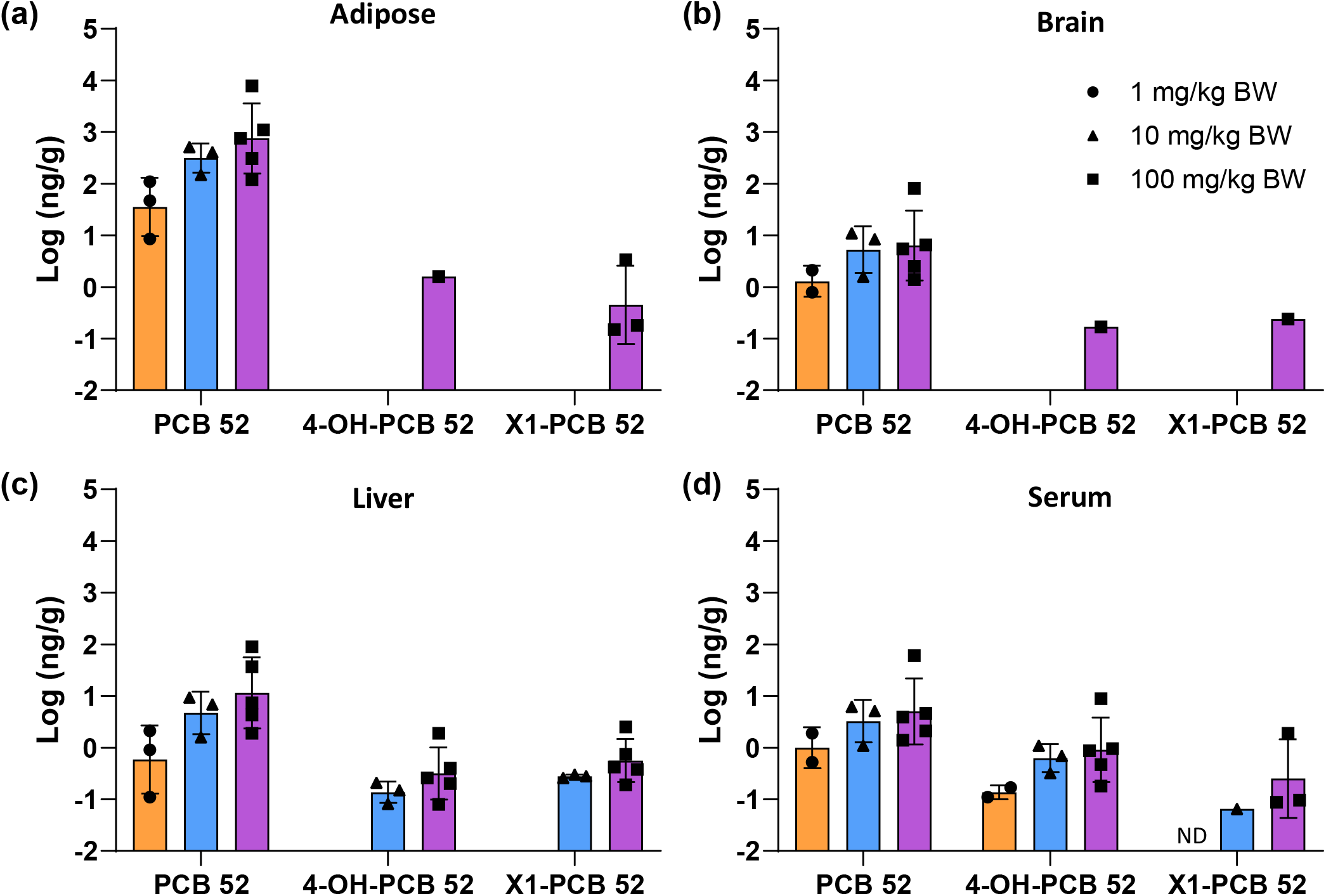
PCB 52 and its mono-hydroxylated metabolite levels were detected in (a) adipose, (b) brain, (c) liver, and (d) serum. Levels expressed as ng/g wet weight are log-transformed to facilitate comparison across tissues with 1 (orange), 10 (blue), and 100 (purple) mg/kg BW, respectively. Data are reported as mean ± standard deviation (N = 3 to 5). ND, not detected.

As ligands for the transport protein transthyretin (TTR) (ATSDR, 2000; Chauhan et al., 2000; Grimm et al., 2015b; 2016), PCBs may cross the blood-brain barrier bound to TTR. However, as lipophilic compounds, PCBs are more likely partition into the brain by passive diffusion. Based on studies of the partitioning of PCBs into lipid bilayers, PCB 52 is expected to cross the blood-brain barrier (Totland et al., 2016). Indeed, PCB 52 was present in the brain (**Fig. 2, Table S7**). PCB 52 levels in the brain were about 50 to 100 times lower than the adipose tissue across all three exposure groups. The extractable lipid content in the mouse brain, expressed as a percent of tissue wet weight, is only 8-fold lower than the lipid content in the adipose tissue (9.3% vs. 72%, respectively) and, thus, cannot explain the drastic difference in PCB 52 levels in both tissues (Wu et al., 2015). Lower PCB levels in the brain than in adipose tissue have been observed in other animal studies and result from the different lipid composition of the brain compared to the adipose tissue (ATSDR, 2000). For example, a composition-based model predicting the PCB disposition in mice poorly approximated the distribution of PCBs from the blood into the brain, most likely because published partition coefficients do not reflect the partitioning of PCBs into brain lipids (Li et al., 2020).

PCB 52 levels in all tissues increased dose-dependent (**Fig. 2, Table S7**). The tissue levels did not reflect the 10-fold differences between doses. The mean PCB 52 levels were approximately 4.0 (serum) to 7.0 (liver) times higher in the tissues from the 10 mg/kg BW than the 1 mg/kg BW dose group. The ratio of the mean PCB 52 tissue levels ranged from only 2.9 (brain) to 5.6 when comparing the 100 mg/kg BW to the 10 mg/kg BW dose group. Other studies have also reported that the PCB tissue ratios between different dose groups do not reflect the fold difference of the dose. For example, female mice exposed orally for 39 days to different doses of PCB 95 showed a 2.5- to 4-fold increase in PCB 95 tissue levels despite a 6-fold difference in the PCB 95 dose (Kania-Korwel et al., 2012). These differences likely reflect the transient induction of hepatic CYPs involved in the oxidative metabolism of PCBs, as suggested by the earlier PCB 95 disposition study in mice. Because CYP2B enzyme activity in the rat liver is induced by PCB 52 exposure (Parkinson et al., 1983), enzymes involved in the metabolism of structurally related PCBs (Lu and Wong, 2011), it is likely that CYPs involved in PCB 52 were transiently induced, thus increasing the elimination of PCB 52 at the higher dose levels. Alternatively, growth dilution could explain the fold differences between exposure groups (Hansen and Welborn, 1977); however, this explanation is unlikely because we did not observe differences in the growth rate (**Fig. 1a**).

### 3.3. Tissue distribution of monohydroxylated PCB 52 metabolites

PCBs with a 2,5-chlorine pattern, like PCB 52, are susceptible to CYP-catalyzed direct electrophilic addition of oxygen to a C-H bond or epoxide formation, ultimately resulting in the formation of isomeric OH-PCBs (Grimm et al., 2015b). Evidence from laboratory and population-based studies demonstrates that these OH-PCBs, like the parent compounds, are also neurotoxic (Grimm et al., 2015b; Ludewig et al., 2008; Tehrani and Van Aken, 2014). OH-PCB isomers may have different modes of action, as shown, for example, by their activity at the level of the ryanodine receptor (Niknam et al., 2013; Sethi et al., 2019). Therefore, we determined the levels of OH-PCBs in all four target tissues using GC-MS/MS (**Fig. 2**).

We detected two metabolites, X1-PCB 52 and 4-OH-PCB 52, in several tissues. The unknown metabolite, X1-PCB 52, is probably 3-OH-PCB 52 (2,2’,5,5’-tetrachlorobiphenyl-3-ol). This assignment is based on the slightly shorter relative retention time of X1-PCB 52 than 4-OH-PCB 52 (relative retention times of 0.788 and 0.80, respectively). Similarly, the retention times of meta vs. para hydroxylated OH-PCBs of multi-*ortho* chlorinated PCB congeners follow the order 1,2-shift << meta < para hydroxylated metabolite (Sundström and Jansson, 1975; Uwimana et al., 2018). Moreover, 3-OH-PCB 52 was formed by phenobarbital-induced rat liver microsomes suggesting a role of CYP2B enzymes in the oxidative metabolism of PCB 52 (Lin et al., 2000). However, studies using an authentic analytical standard are needed to confirm this tentative identification of X1-PCB 52. Levels of OH-PCBs across all tissues were lower compared to PCB 52 levels, consistent with an earlier disposition study with other PCB congeners in rats (Dhakal et al., 2014).

X1-PCB 52 was present in the adipose tissue from the high-dose group but below the LOD in all other exposure groups (**Fig. 2**). 4-OH-PCB 52 was detected only in adipose tissue from one animal in the high-dose group. Previous work from our group also did not detect OH-PCBs in the adipose of PCB-exposed mice (Kania-Korwel et al., 2015; Kania-Korwel et al., 2012; Wu et al., 2020). In contrast, other PCB metabolites have been discovered in other organisms, such as MeSO_2_-PCB metabolites in the blubber of gray seals (Jensen et al., 1979). MeSO_2_-PCBs were also some of the first PCB metabolites identified in human adipose tissue from a woman exposed via the occupational handling of PCB-containing capacitors (Yoshida and Nakamura, 1977). Primarily higher chlorinated PCB metabolites have been reported previously in human liver and adipose samples (Guvenius et al., 2002). These results suggest that some PCB metabolites, including OH-PCBs, can be present at low levels in the adipose and, in addition to the parent PCBs, may contribute to toxic outcomes. In a recent study in which human preadipocytes were treated with PCB 52 or 4-OH-PCB 52 in vitro, we found that 4-OH-PCB 52 caused both different and considerably more changes in the number of differentially expressed genes as compared to PCB 52 (Gourronc et al., 2023), suggesting that OH-PCBs are more biologically active and potentially more toxic to adipose than the parental PCB. Similarly, metabolites of other environmental contaminants, such as polycyclic aromatic hydrocarbons (PAHs), can be more toxic than the parent compounds (Lam et al., 2018).

In this study, 4-OH-PCB 52 and X1-PCB 52 were detected in the livers of animals from the 10 mg/kg BW (mean levels of 0.15 and 0.28 ng/g, respectively) and 100 mg/kg BW (mean levels of 0.57 and 0.85 ng/g, respectively) exposure groups. OH-PCBs have been previously identified in the livers of mice and their offspring orally exposed subacutely and subchronically to PCB 95, a PCB congener structurally related to PCB 52 (Kania-Korwel et al., 2012; Kania-Korwel et al., 2017). Interestingly, OH-PCB 95 metabolite levels were 4 to 900-fold higher in these earlier studies than in this study. These differences in the OH-PCB levels are likely due to the different routes of exposure, the dosing paradigms, and structural differences. In human liver samples, OH-PCB levels (∑OH-PCBs between 7 to 175 ng/g lipids) were about 10 to 1000-fold higher than in the present study (Guvenius et al., 2002).

We found estimated mean X1-PCB 52 levels in the serum of 0.07 ng/kg BW for the 10 mg/kg BW and 0.69 ng/g for the 100 mg/kg BW exposure group. In addition, 4-OH-PCB was present in the serum of animals with mean levels of 0.14, 0.70, and 2.27 ng/g for animals exposed to 1, 10, or 100 mg/kg BW PCB 52, respectively (**Table S7**). Serum PCB metabolites are routinely assessed in rodents (Stamou et al., 2015; Zhang et al., 2021), wildlife (Quinete et al., 2014), and humans (Koh et al., 2016a; Koh et al., 2016b; Quinete et al., 2017; Quinete et al., 2014). Another study with rural and urban participants identified 58 OH-PCBs at mean levels between 0.05 to 0.08 ng/g in the serum of both adults and adolescents (Koh et al., 2016a; Koh et al., 2016b). Previously in humans, 38 OH-PCB metabolites have been reported in plasma, many of which have five or more chlorine substitutes (Hovander et al., 2002). More work reported total median levels of OH-PCBs in different human populations between 0.0002 and 1.6 ng/g wet weight in serum/plasma (Quinete et al., 2014). In addition, lower chlorine PCB metabolites, such as OH-PCB 28 metabolites, are present in human plasma, with total OH-PCB levels ranging from 0.03 to 27.7 ng/mL plasma (Quinete et al., 2017). These total OH-PCB levels are comparable to the OH-PCB 52 levels detected in the serum in this study (**Table S7**); however, OH-PCB 52 metabolites have not been reported in human serum to date, most likely because the corresponding analytical standards are not commercially available.

X1-PCB 52 and 4-OH-PCB 52 were also observed in the brain of one animal. The presence of hydroxylated PCB 52 metabolites in the brain has not been reported previously. However, a single cysteinyl adduct of a 1,2-benzoquinone of PCB 52 was detected in the brain of rats exposed to PCB 52 (Lin et al., 2000). Several other studies have reported the presence of OH-PCBs in the mammalian brain. One study identified radiolabeled 4-OH-PCB 107 in the forebrain and cerebellum of Wistar dams and their offspring (Meerts et al., 2002). Our group has identified OH-PCB 95 metabolites at low levels in the brain (11 to 14 ng/g tissue across doses from 0.1 to 6.0 mg/kg body weight per day) of C57Bl/6 mice exposed subchronically to different PCB 95 doses (Kania-Korwel et al., 2012). Several OH-PCBs were present in the brains of dogs exposed intraperitoneally to a synthetic PCB mixture (Nomiyama et al., 2019). In the Japanese macaque (*Macaca fuscata*), OH-PCBs were present in the fetal brain (Nomiyama et al., 2020). Another study did not detect OH-PCBs in the mouse brain, possibly because of the comparatively high detection limit of the analytical method used in these studies. For example, no OH-PCB 136 metabolites were detected in the brain of mice exposed to PCB 136 (Wu et al., 2015). Recently, OH-PCB metabolites of several lower chlorinated PCBs were detected in postmortem human brain tissue (Li et al., 2022).

### 3.4. Nt-LCMS analysis of PCB 52 conjugates

Complex mixtures of PCB 52 metabolites, including PCB sulfate and other conjugates, may be formed in PCB 52 exposed rats. These metabolites, for example, cysteinyl adducts of PCB quinone metabolites, may also be present in the brain and can be neurotoxic (Grimm et al., 2015b). Because the metabolism of PCB 52 in rats has not been fully characterized, we employed an Nt-LCMS method to semiquantitatively identify PCB 52 metabolites present in target tissues and elucidate the metabolic pathway of PCB 52 in rats. This Nt-LCMS method has been used previously to characterize PCB metabolite profiles in cell culture (Zhang et al., 2020) and animal models (Li et al., 2021). The Nt-LCMS analyses of liver and serum samples revealed several PCB 52 metabolites (**Table 1**). Several metabolites were only detected in one animal exposed to the high PCB 52 dose. This animal also displayed higher PCB 52 and OH-PCB 52 levels than animals from the same exposure group. Metabolites identified in both tissues included mono-hydroxylated PCB 52 metabolites and the corresponding sulfates. We also observed several methoxylated and dechlorinated metabolites, consistent with previous PCB metabolism studies in cells in culture (Zhang et al., 2020) and in the urine of rabbits exposed to PCB 153 through diet (Hutzinger et al., 1974), the feces of mice, rats, and quails after consumption of PCB 95 (Sundström and Jansson, 1975), or through metabolism of PCB 3 following intraperitoneal exposure (Safe et al., 1975). Importantly, no PCB 52 conjugates were detected in the brain and adipose tissue, including the cysteinyl adduct of a PCB 52 quinone metabolite reported previously (Lin et al., 2000). It is important to emphasize that, due to a 1,2 shift of PCB 52 epoxide intermediates, metabolites may not have retained the 2,2’,5,5’-tetrachloro substitution pattern (Guroff et al., 1967).

**Table 1.** Several PCB 52 metabolite classes were detected by LC-Orbitrap MS in tissues from female rats exposed to 100 mg/kg BW PCB 52 via I.P. injection.

| Class No | Metabolites<sup>a</sup> | Retention time (min) | Formula | Normalized intensity<sup>b</sup> | [M-H]<sup>-</sup> |  | Accurate mass difference, ppm<sup>c</sup> | Confidence level |
| --- | --- | --- | --- | --- | --- | --- | --- | --- |
|  |  |  |  |  | Calculated (Da) | Measured (Da) |  |  |
| Liver |  |  |  |  |  |  |  |  |
| 1.1 |  | 7.71 | C<sub>12</sub>H<sub>5</sub>OC<sub>14</sub> | 0.18 ± 0.20 | 304.91000 | 304.90991 | -0.2952 | 3 |
| 1.2 | Tetra-CB sulfate | 6.34 | C<sub>12</sub>H<sub>5</sub>Cl<sub>4</sub>O<sub>4</sub>S | 0.22 ± 0.34 | 384.86681 | 384.86694 | 0.3378 | 2 |
|  |  | 6.41 |  | 0.19 ± 0.29 | 384.86681 | 384.86691 | 0.2598 | 3 |
| 1.3 | diOH-Tetra-CB | 6.63 | C<sub>12</sub>H<sub>5</sub>Cl<sub>4</sub>O<sub>2</sub> | 0.29 | 320.90491 | 320.90485 | -0.1870 | 3 |
| 1.4 | OH-Tetra-CB sulfate | 5.42 | C<sub>12</sub>H<sub>5</sub>O<sub>5</sub>Cl<sub>4</sub>S | 0.35 ± 0.54 | 400.86173 | 400.86176 | 0.0748 | 2 |
|  |  | 5.62 |  | 0.12 ± 0.13 | 400.86173 | 400.86172 | -0.0249 | 2 |
|  |  | 6.77 |  | 0.74 | 400.86173 | 400.86145 | -0.6985 | 2 |
| 1.5 | MeO-OH-Tetra-CB | 7.86 | C<sub>13</sub>H<sub>7</sub>O<sub>3</sub>Cl<sub>4</sub> | 0.09 | 334.92056 | 334.9205 | -0.1791 | 3 |
| 1.6 | MeO-OH-Tetra-CB sulfate | 5.28 | C<sub>13</sub>H<sub>7</sub>O<sub>6</sub>Cl<sub>4</sub>S | 0.05 | 430.87229 | 430.87070 | -3.6902 | 2 |
|  |  | 5.47 |  | 0.01 | 430.87229 | 430.87235 | 0.1393 | 2 |
|  |  | 5.57 |  | 0.02 | 430.87229 | 430.87238 | 0.2089 | 2 |
|  |  | 5.67 |  | 0.01 | 430.87229 | 430.87188 | -0.9516 | 2 |
| 2.1 | OH-Tri-CB sulfate | 6.01 | C<sub>12</sub>H<sub>6</sub>C<sub>13</sub>O<sub>5</sub>S | 0.12 | 366.90070 | 366.90070 | 0.0000 | 2 |
| 2.2 | MeO-Tri-CB sulfate | 5.98 | C<sub>13</sub>H<sub>8</sub>Cl<sub>3</sub>O<sub>5</sub>S | 0.04 | 380.91635 | 380.91669 | 0.8926 | 2 |
| 2.3 | MeO-diOH-Tri-CB | 6.49 | C<sub>13</sub>H<sub>8</sub>Cl<sub>3</sub>O<sub>3</sub> | 0.02 | 316.95445 | 316.95352 | -2.9342 | 3 |
| 2.4 | MeO-OH-Tri-CB sulfate | 5.47 | C<sub>13</sub>H<sub>8</sub>Cl<sub>3</sub>O<sub>6</sub>S | 0.42± 0.20 | 396.91127 | 396.91125 | -0.0504 | 2 |
| Serum |  |  |  |  |  |  |  |  |
| 3.1 |  | 7.70 | C<sub>12</sub>H<sub>5</sub>OC<sub>14</sub> | 1.10 | 304.91000 | 304.90942 | -1.9022 | 3 |
| 3.2 | Tetra-CB sulfate | 6.34 | C<sub>12</sub>H<sub>5</sub>Cl<sub>4</sub>O<sub>4</sub>S | 1.04 | 384.86681 | 384.86713 | 0.8315 | 2 |
| 3.3 | OH-Tetra-CB sulfate<sup>a</sup> | 6.41 | C<sub>12</sub>H<sub>5</sub>O<sub>5</sub>Cl<sub>4</sub>S | 1.47 ± 2.07 | 400.86173 | 400.86133 | -0.9979 | 2 |
|  |  | 6.63 |  | 0.63 ± 0.97 | 400.86173 | 400.86066 | -2.6693 | 2 |
| 3.4 | MeO-OH-Tetra-CB | 5.42 | C<sub>13</sub>H<sub>7</sub>O<sub>3</sub>Cl<sub>4</sub> | 0.79 | 334.92056 | 334.91989 | -2.0005 | 3 |
| 3.5 | MeO-OH-Tetra-CB sulfate | 5.26 | C<sub>13</sub>H<sub>7</sub>O<sub>6</sub>Cl<sub>4</sub>S | 0.07 | 430.87229 | 430.87260 | 0.7195 | 2 |
|  |  | 5.47 |  | 0.02 | 430.87229 | - | - | 2 |
| 4.1 | OH-Tri-CB sulfate | 6.02 | C<sub>12</sub>H<sub>6</sub>C<sub>13</sub>O<sub>5</sub>S | 0.03 | 366.90070 | 366.90036 | -0.9267 | 2 |
| 4.2 | MeO-OH-Tri-CB sulfate | 5.06 | C<sub>13</sub>H<sub>8</sub>C<sub>13</sub>O<sub>6</sub>S | 0.10 | 396.91127 | 396.90991 | -3.4265 | 2 |
|  |  | 5.47 |  | 0.37 | 396.91127 | 396.91052 | -1.8896 | 2 |

<sup>a</sup> Metabolite also detected in 10 mg/kg PCB 52 exposed animals
<sup>b</sup> Some metabolites were only detected in one animal
<sup>c</sup> parts per million, ppm = (measured -calculated)/calculated\*1000000
CB, chlorobiphenyl

#### 3.4.1. Identification of PCB 52 metabolites in the liver by Nt-LCMS

The Nt-LCMS analysis revealed the presence of several PCB metabolites in the liver (**Fig. 3B, Table 1**; see also **Figs. S3 to S12**). For example, two PCB 52 sulfate compounds were detected in the liver, consistent with the presence of two OH-PCB isomers in the GC-MS/MS analysis. OH-PCBs undergo sulfation to PCB sulfates by rat SULT in gene family 1, more specifically SULT1A1 (Liu et al., 2006; Liu et al., 2009); however, the formation of PCB 52 sulfates in rats *in vivo* has not been reported previously. Moreover, other studies have reported the presence of other PCB sulfates in the liver of rodents exposed to lower chlorinated PCBs (Dhakal et al., 2014; Zhang et al., 2021). The Nt-LCMS analysis also identified one dihydroxylated OH-PCB metabolite in the liver of PCB 52 exposed rats, consistent with earlier *in vitro* and *in vivo* studies reporting the formation of 2,2’,5,5’-tetrachlorobiphenyl-3,4-diol. Dihydroxylated PCB metabolites can be formed from the monohydroxylated metabolites in secondary oxidation reactions (Grimm et al., 2015b). Our analysis detected several di-OH-PCB-derived metabolites, including three OH-PCB sulfates and one tetrachlorinated MeO-OH-PCB metabolite. Finally, we tentatively identified four tetrachlorinated MeO-OH-PCB sulfates, metabolites derived from tri-hydroxylated PCBs (**Fig. S8**). These metabolites of PCB 52 are in agreement with earlier metabolism studies with LC-PCBs *in vitro* (McLean et al., 1996), cells in culture (Zhang et al., 2020), in rats *in vivo* (Grimm et al., 2015b; Sundström and Jansson, 1975), and humans (Zhang et al., 2022b). The formation of di- and tri-hydroxylated PCB 52 metabolites and their conjugates is a concern because these hydroxylated metabolites can cause toxic effects through intracellular oxidative stress mechanisms (Song et al., 2008).

**Fig. 3.**
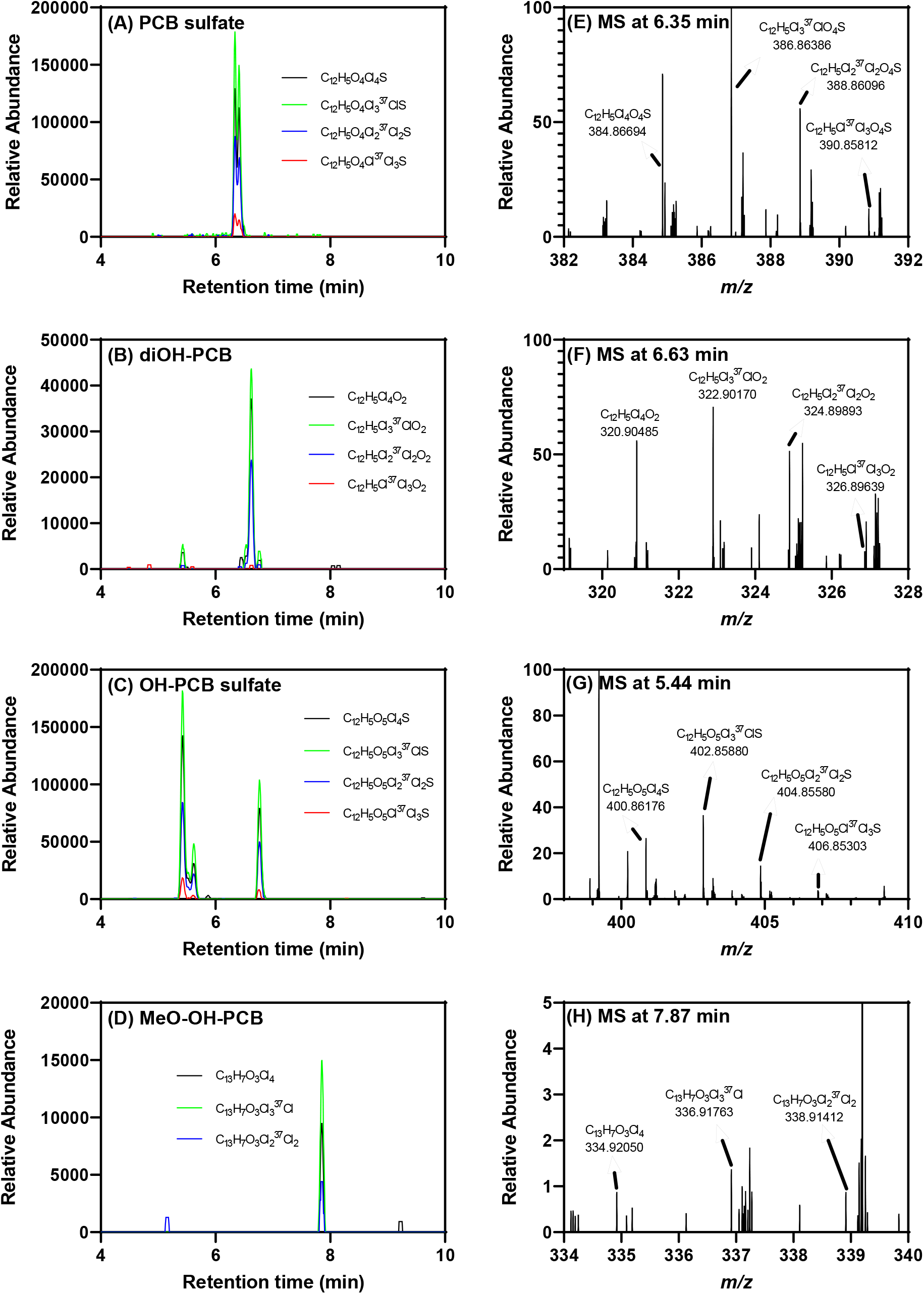
Several tetrachlorinated PCB metabolites were detected by LC-Orbitrap MS in the liver of a rat exposed to 100 mg/kg BW of PCB 52, including (A) two PCB sulfates ([C_12_H_5_Cl_4_O_4_S]^-^, *m/z* 384.86681), (B) one diOH-PCB ([C_12_H_5_Cl_4_O_2_]^-^, *m/z* 320.90491), (C) three OH-PCB sulfate ([C_12_H_5_O_5_Cl_4_S]^-^, *m/z* 400.86173), and (D) one MeO-OH-PCB ([C_13_H_7_O_3_Cl_4_]^-^, *m/z* 334.92056) peaks. The presence of the (E) PCB sulfate at 6.35 min, (F) diOH-PCB at 6.63 min, (G) OH-PCB sulfate at 5.44 min, and (H) MeO-OH-PCB at 7.87 min was confirmed based on the accurate masses and isotopic pattern of isotope ions corresponding to a tetrachlorinated PCB metabolite. The LC-Orbitrap MS analysis was performed in the negative polarity mode. *m/z* values are reported for the monoisotopic ion. More information, including the MS data of other tetrachlorinated metabolites detected in the same extract, are included in the supporting information.

Along with tetrachlorinated metabolites, we also observed trichlorinated PCB metabolites in the liver of PCB 52 exposed rats, including one OH-PCB sulfate and one MeO-OH-PCB sulfate (**Fig. 4**; see also **Figs. S9 to S12**). This observation indicates a metabolism pathway that results in the dehalogenation of the biphenyl moiety. The loss of chlorine from metabolizing PCB 95 was observed in feces extracts of rats (Sundström and Jansson, 1975). In addition, we tentatively identified one trichlorinated MeO-PCB sulfate (**Fig. S10**) and one trichlorinated MeO-diOH-PCB (**Fig. S11**). Similar dechlorinated PCB 2 and PCB 11 metabolites were also formed by HepG2 cells, a human hepatocarcinoma cell line. Metabolism studies with PCB 2 and selected OH-PCB 2 metabolites demonstrated that these diOH-PCB-derived metabolites are formed by the displacement of a Cl group adjacent to an OH-group by OH (Zhang et al., 2022a).

**Fig. 4.**
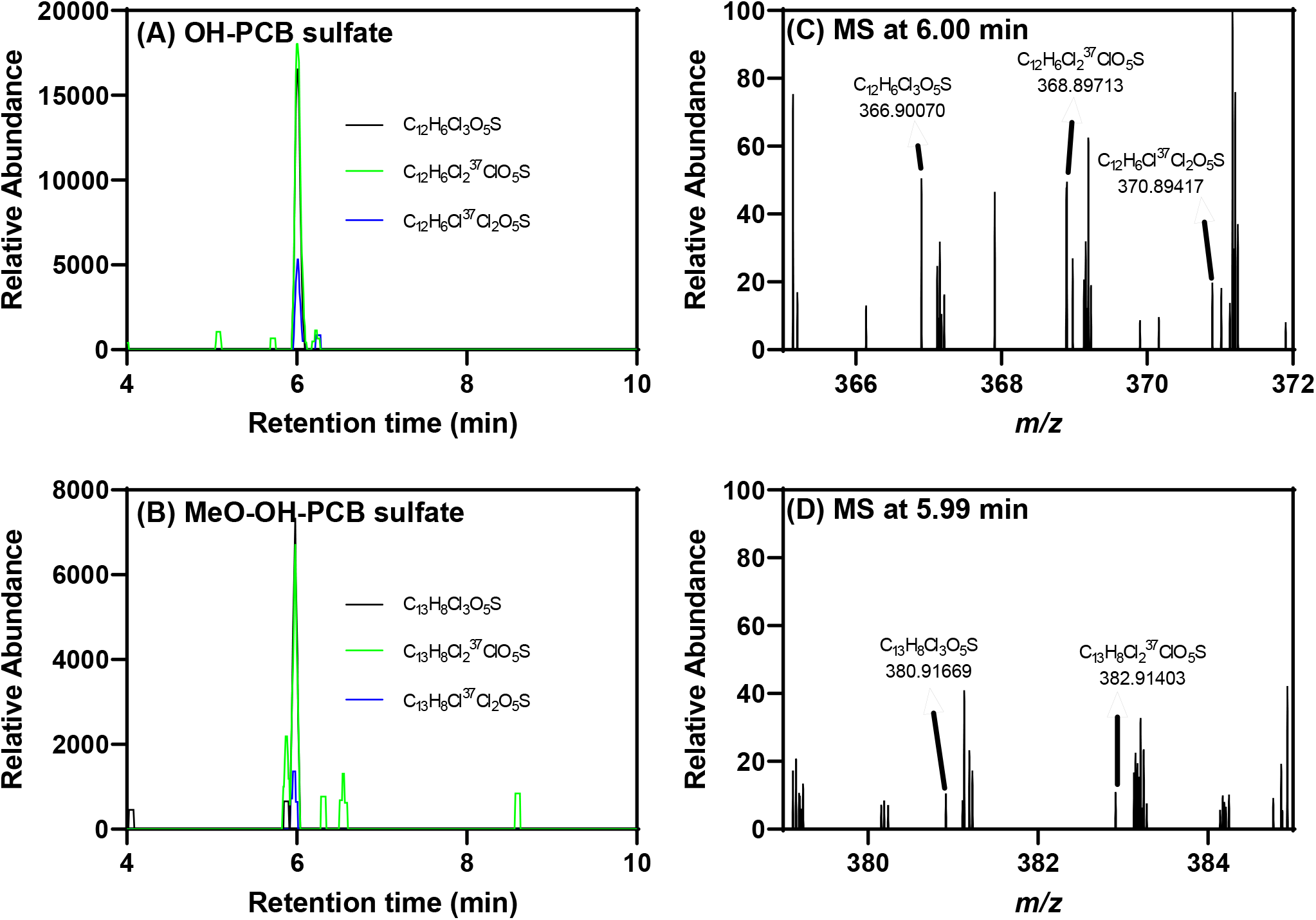
Select trichlorinated PCB metabolites detected by LC-Orbitrap MS in the liver of PCB 52 exposed rats including (A) one OH-PCB sulfate ([C_12_H_6_C_l3_O_5_S]^-^, *m/z* 366.90070), (B) one MeO-OH-PCB sulfate ([C_13_H_8_Cl_3_O_6_S]^-^, *m/z* 396.91127) peaks. The presence of the (C) trichlorinated OH-PCB sulfate at 6.00 min and (D) trichlorinated MeO-OH-PCB sulfate at 5.99 min was confirmed based on the accurate masses and isotopic pattern of isotope ions corresponding to a trichlorinated PCB metabolite. The LC-Orbitrap MS analysis was performed in the negative polarity mode. More information found in the supporting information.

#### 3.4.2. Identification of PCB 52 metabolites in the serum by Nt-LCMS

The Nt-LCMS analysis identified several compounds in the serum of animals exposed to PCB 52, including two OH-PCB 52 sulfates and one MeO-OH-PCB 52 (**Fig. 5**; see also **Figs. S13 to S18**). In addition, one PCB 52 sulfate and one MeO-OH-PCB 52 sulfate were tentatively identified by Nt-LCMS. The PCB sulfate formed from 4-OH-PCB 52 has been detected in human serum (Zhang et al., 2022b). Like the dechlorinated PCB metabolites identified in the liver, the serum also contained two trichlorinated MeO-OH-PCB sulfates, providing further evidence that dechlorinated PCB metabolites are formed in rats *in vivo* (Hutzinger et al., 1974; Sundström and Jansson, 1975). Additional preclinical studies and human biomonitoring studies are needed to confirm the formation of dechlorinated PCB metabolites in humans.

**Fig. 5.**
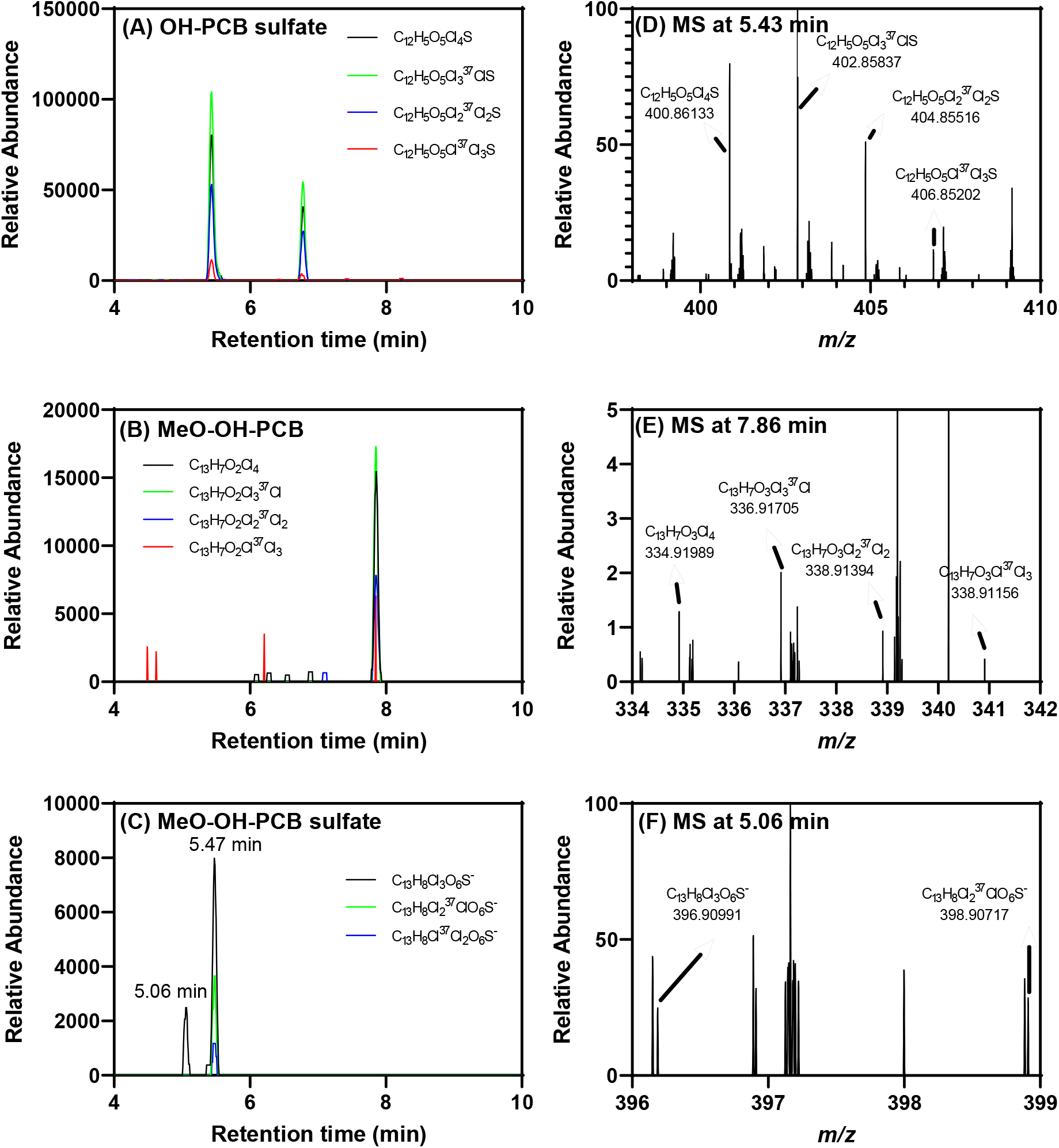
Select PCB metabolites were detected by LC-Orbitrap MS in the serum of PCB 52 exposed rats, including (A) two tetrachlorinated OH-PCB sulfate ([C_12_H_5_O_5_Cl_4_S]^-^, *m/z* 400.86173), (B) one tetrachlorinated MeO-OH-PCB ([C_13_H_7_O_3_Cl_4_]^-^, *m/z* 334.92056), and (C) two trichlorinated MeO-OH-PCB sulfates ([C_13_H_8_C_l3_O_6_S]^-^, *m/z* 396.90991) peaks. The presence of the (D) OH-PCB sulfate at 5.43 min, (E) MeO-OH-PCB at 7.86 min, and (F) trichlorinated MeO-OH-PCB sulfate at 5.06 min was confirmed based on the accurate masses and isotopic pattern of isotope ions corresponding to a tetrachlorinated PCB metabolite for the monoisotopic ion). The accurate masses of two high-abundance isotope ions at (F) 5.06 match the theoretical accurate mass and isotopic pattern of a trichlorinated compound. The LC-Orbitrap MS analysis was performed in the negative polarity mode, as described in the Supporting Information. More information in the supporting information.

#### 3.4.3 Metabolism pathway of PCB 52 in rats

Based on the Nt-LCMS analysis results, we propose the metabolic pathway for PCB 52 in rats shown in **Fig. 6**. Briefly, PCB 52 is oxidized to mono-hydroxylated OH-PCB metabolites by CYPs. The OH-PCB metabolites are metabolized to the corresponding sulfates by sulfotransferase (SULT) or oxidized by CYPs to di- and tri-hydroxylated PCB metabolites. These metabolites undergo further SULT or catechol-O-methyltransferase (COMT) catalyzed biotransformation to diverse sulfated and methoxylated PCB 52 metabolites. Also, OH-PCB metabolites undergo chlorine displacement to form dechlorinated dihydroxylated PCB metabolites that undergo further biotransformation to sulfate conjugates and methoxylated PCBs. It is also possible that OH-PCB sulfate metabolites are formed via the oxidation of PCB 52 sulfates. Consistent with this metabolism pathway, 4-PCB 11 sulfate is metabolized to further biotransformation products in rats exposed intravenously to 4-PCB 11 sulfate (Grimm et al., 2015a). Several metabolites, including MeO-PCBs and di-OH-PCBs, may also be formed by deconjugation reactions. For example, human hepatic microsomal sulfatases catalyze the deconjugation of PCB sulfates to the corresponding OH-PCBs (Duffel et al., 2021). Overall, the PCB 52 metabolism pathway outlined in **Fig. 6** is consistent with published *in vitro* and *in vivo* studies (Grimm et al., 2015b). However, as reported previously, we did not observe the formation of glutathione or cysteine adducts of PCB 52 quinones (Lin et al., 2000). These differences may be due to the different routes of exposure (intraperitoneal vs. oral gavage) or the different analytical methods.

**Fig. 6.**
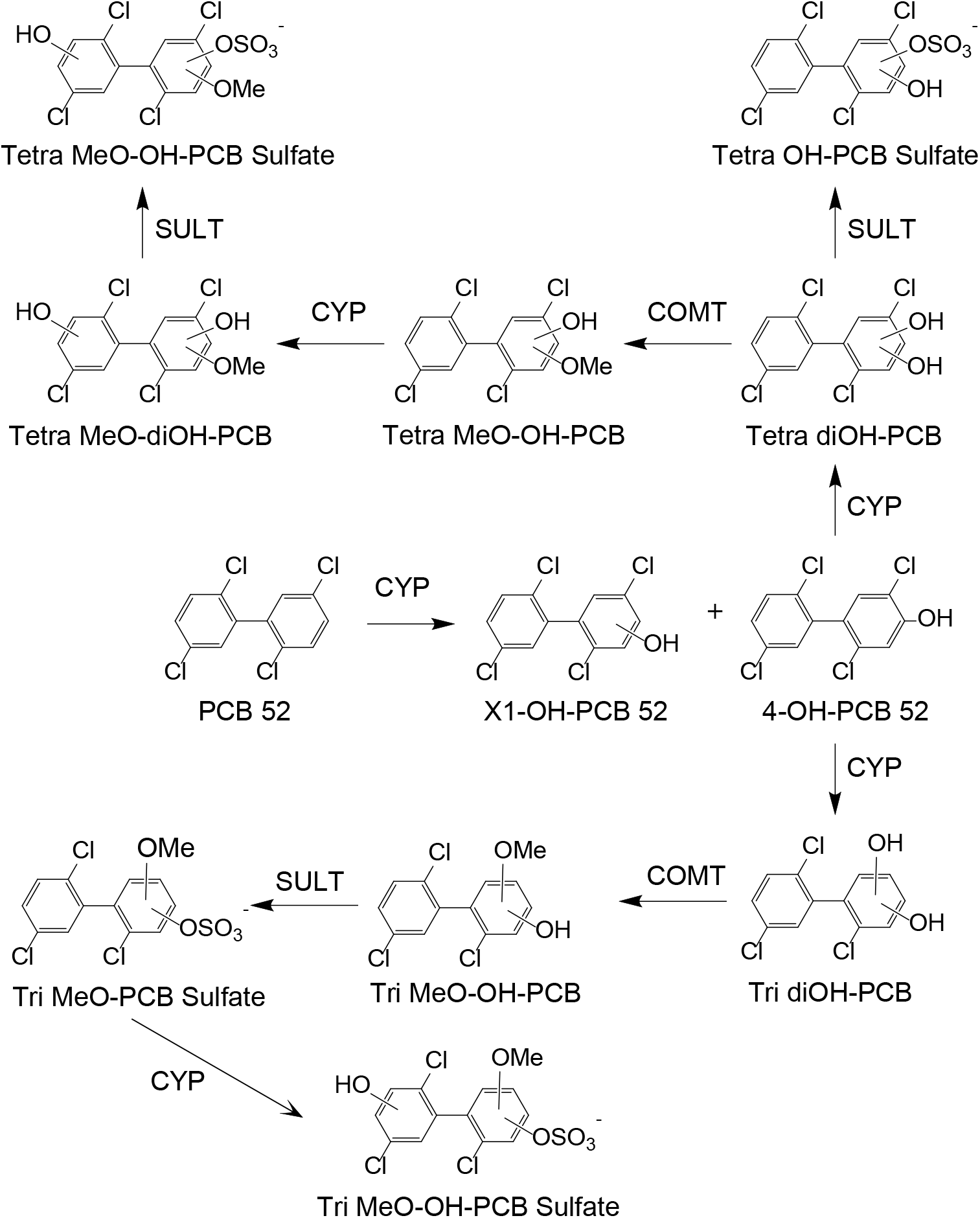
Proposed PCB 52 metabolic pathway in rats based on the metabolites detected in the targeted GC-MS/MS and the Nt-LCMS analyses, see text for details. CYP, cytochrome P450 enzyme; SULT, sulfotransferase; COMT, catechol-O-methyltransferase.

### 3.5. Changes in serum metabolome of PCB 52 exposed female rats

We investigated changes in endogenous metabolic pathways in the serum of female Sprague Dawley rats following exposure to PCB 52 to assess if PCB 52 and metabolite levels are potentially associated with changes in specific metabolic pathways. The targeted metabolomic analysis identified only 9 significantly altered cell metabolites out of the 81 metabolites included in the data analysis (**Fig. 7)**, thus making it challenging to study associations between PCB levels and metabolomic outcomes. These results suggest that PCB 52 may affect some metabolic functions *in vivo*. However, because the metabolomics analyses were performed in samples taken three weeks after PCB 52 exposure, further research is necessary to fully understand the effects of PCB 52 exposure on serum metabolome over time and after repeated exposure.

**Fig. 7.**
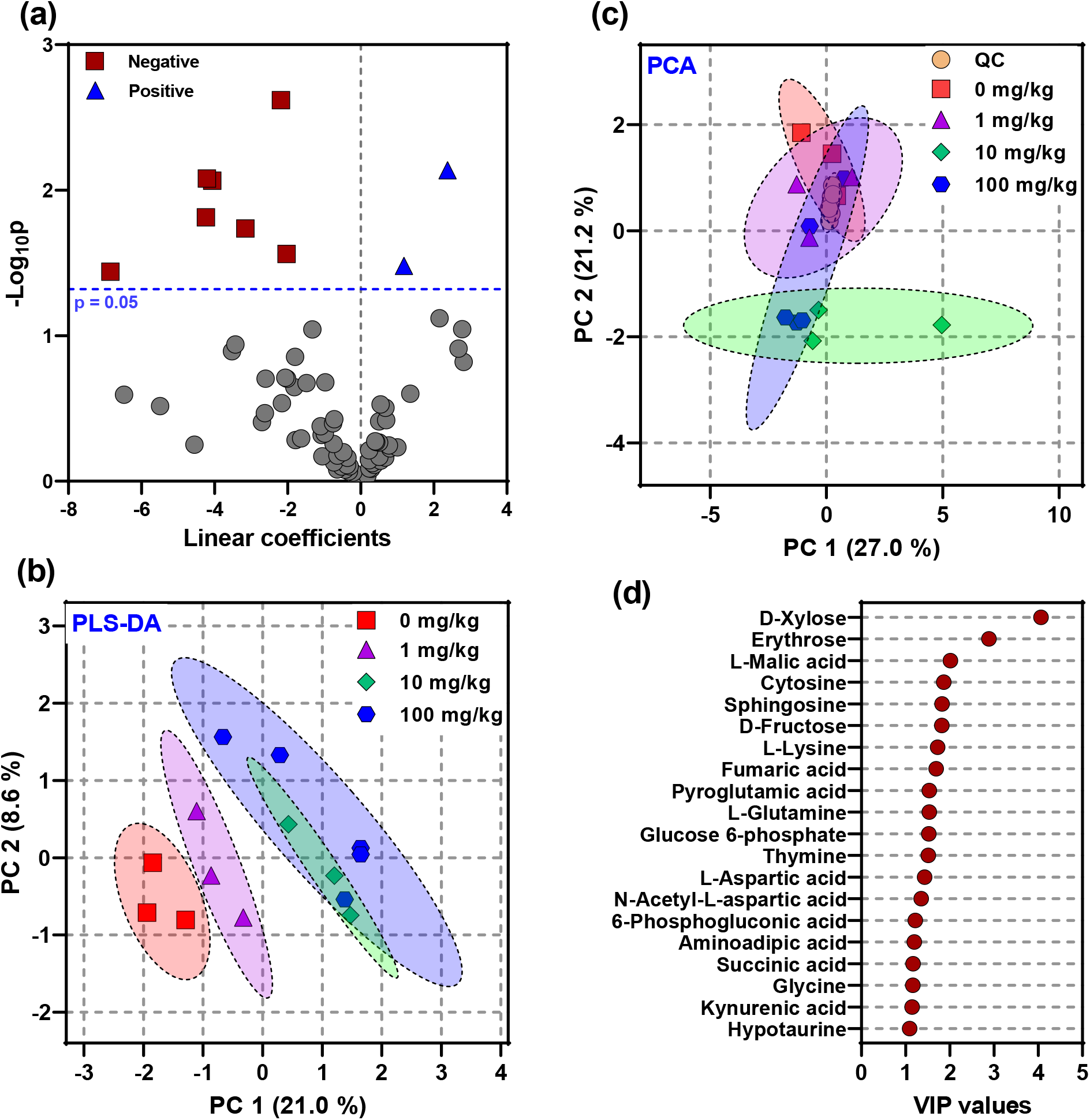
Targeted metabolomic analysis suggested a small number of serum metabolites associated with acute IP exposure to PCB 52 doses ranging from 0 to 100 mg/kg body weight. (a) Volcano plot with p-values from an univariate analysis identified seven and two metabolites negatively or positively associated with PCB 52 exposure, respectively. For bar plots of these significantly associated metabolites, see **Fig. 8**. Multivariate analysis, including (b) principal component analysis (PCA) and (c) partial least squares-discriminant analysis (PLS-DA), revealed reasonable data variability and potential metabolites that drive the dose-dependent clustering, as indicated by the (d) variable importance in projection (VIP) values in the PLS-DA. For information regarding the metabolomic analysis, see Materials and Methods.

L-cysteine, cytosine, sphingosine, orotic acid, and erythrose levels decreased dose-dependent following PCB 52 exposure. A decrease in L-cysteine levels has been implicated in male reproductive maladaptive responses following PCB exposure in rats (Mega et al., 2022). The reduction in sphingosine suggests alterations in gap-junction intracellular communication via changes in sphingolipid levels. PCB-mediated effects on sphingolipid levels in the liver decrease gap-junction intracellular communication, a phenomenon implicated in tumor regulation (Pierucci et al., 2017). Finally, the PCB mixture Phenoclor DP6 reduced the incorporation of the orotic acid-derived nucleotides thymine and cytosine into the nuclei of liver cells in PCB-exposed rats (Narbonne, 1979). The effects of PCB exposure on erythrose have not been reported previously. However, a recent study reported that PCBs increased levels of an erythrose phosphate, D-erythrose-4-phosphate, in the serum of cats, potentially affecting the pentose pathway (Nomiyama et al., 2022).

Glycine and L-histidine levels showed a dose-dependent increase following PCB 52 exposure (**Fig. 8**). Effects of PCB 52 and structurally related PCBs on these two metabolites *in vivo* have not been reported previously. Furthermore, PCB 52 significantly affected thymine and linoleic acid levels in the serum of female rats; however, no dose-response relationship was observed for either metabolite. The observation that PCB 52 exposure affected linoleic acid metabolism is not surprising because PCB congeners with multiple *ortho* chlorine substituents are known to significantly change the fatty acid composition, particularly in the rat liver (Azaïs-Braesco et al., 1990; Matsusue et al., 1997).

**Fig. 8.**
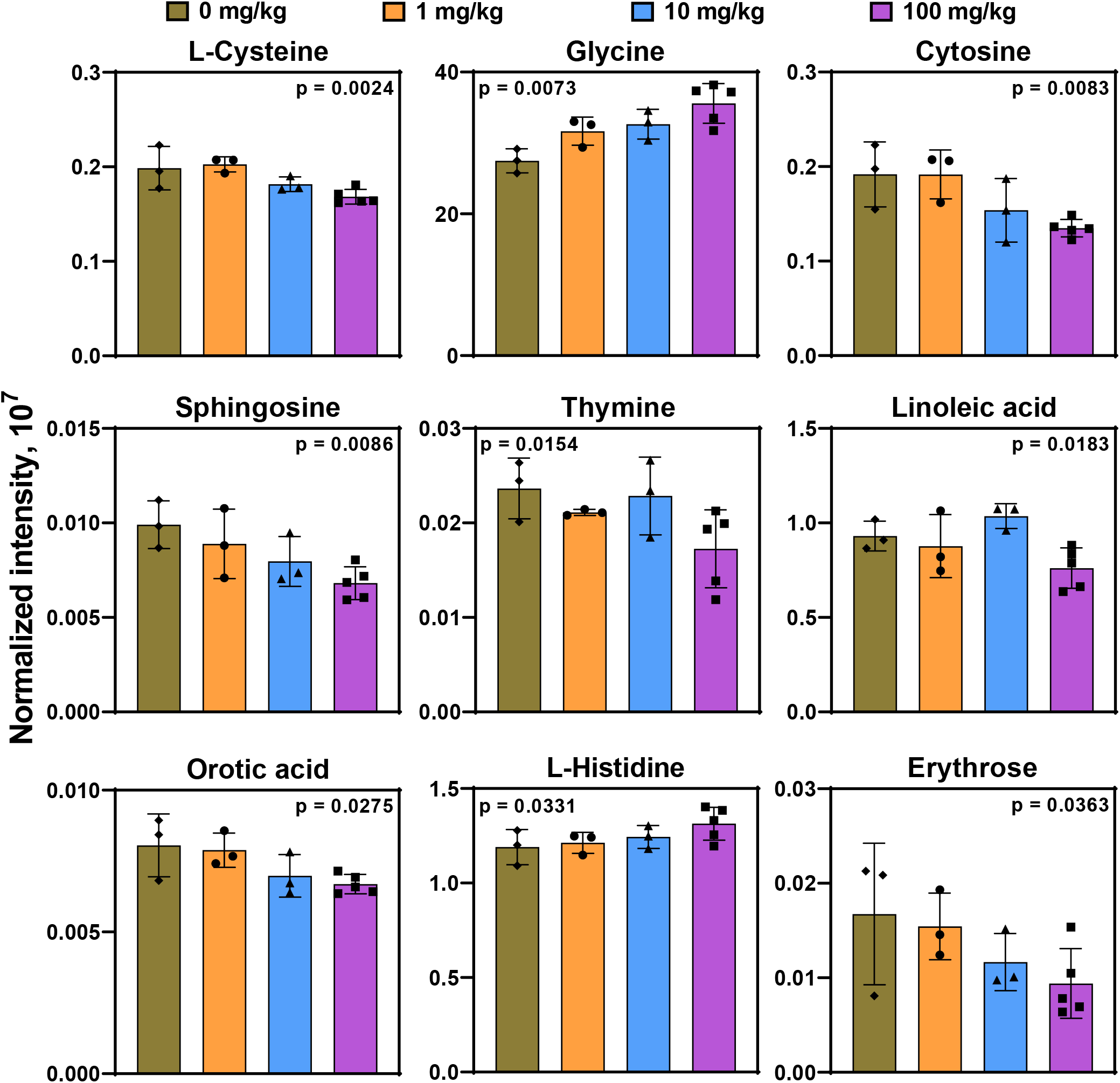
Univariate analysis with a linear regression model selected nine dose-dependent serum metabolites in rats exposed via IP injection to PCB 52 with doses ranging from 0 to 100 mg/kg body weight. Data are reported as mean ± standard deviation. Intensities were normalized with the total intensity of each sample.

### 3.6. Changes in striatal dopamine and metabolites after PCB 52 exposure in female rats

Several individual PCB congeners and complex PCB mixtures influence the dopaminergic system *in vivo*. Briefly, studies with rat brain synaptosomes demonstrate that *ortho*-substituted PCBs significantly inhibit dopamine uptake into vesicles (Mariussen et al., 2001; Mariussen and Fonnum, 2001). Studies in cells in culture reported conflicting effects of individual, *ortho*-chlorinated PCB congeners on dopamine levels, consistent with different, structure-dependent mechanisms by which PCBs affect cellular dopamine metabolism (Enayah et al., 2018; Uwimana et al., 2019). *In vivo*, dopamine levels were significantly decreased in several brain regions of pig-tailed macaques following oral exposure to Aroclor 1016 or Aroclor 1260 (Seegal et al., 1994; Seegal et al., 1990). Another study reported decreased dopamine transporters (DAT) protein levels and dopamine uptake in mice exposed to Aroclor 1254 and Aroclor 1260 (Caudle et al., 2006). In contrast, a 4-week oral exposure in rats to a PCB mixture containing PCB 52 increased striatal dopamine levels 2 weeks post-exposure (Lee et al., 2012).

Here, we investigated if PCB 52 exposure affects the levels of dopamine and its metabolites, DOPAL and DOPAC. We observed no significant differences in dopamine, DOPAL, or DOPAC levels in the striatum of PCB 52 exposed animals (**Fig. 9**). No changes in either dopamine or DOPAC levels were reported by a study that exposed rats for 91 days to an inhaled mixture of PCBs (Wang et al., 2022). It is possible that the effects of PCB 52 on dopamine homeostasis in the rat brain were transient and, therefore, could not be detected three weeks after PCB 52 exposure. Moreover, PCBs or their metabolites may differentially affect the expression of dopaminergic genes, such as the dopamine receptor, in dopaminergic brain regions (Lesmana et al., 2014; Liberman et al., 2020), thus masking changes in dopamine homeostasis in our study. Finally, the metabolism of PCB 52 to hydroxylated metabolites may be necessary to alter dopamine levels in the female rat brain.

**Fig. 9.**
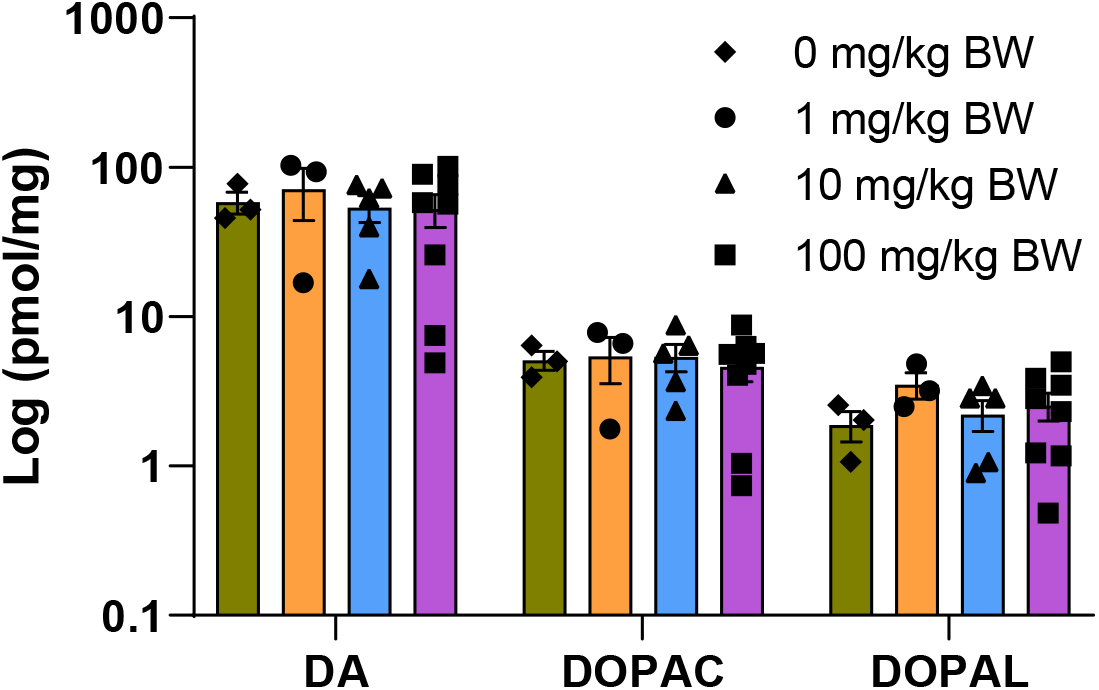
Dopamine and metabolite concentrations from the striatum of PCB 52 exposed female rats. Concentrations are plotted on a log_10_ scale and reported in pmol per mg of striatal tissue processed. Dopamine (DA), 3,4-Dihydroxyphenylacetic acid (DOPAC), or 3,4-dihydroxyphenylacetaldehyde (DOPAL) concentrations were not significantly different 3 weeks post IP injection of different doses of PCB 52. Group means were compared using two-way ANOVA followed by Tukey’s pairwise comparisons. A p-value < 0.05 was considered statistically significant.

## 4. Conclusions

Because only limited studies of the disposition and metabolomic effects of PCB 52 are available, this exploratory study characterized the disposition of PCB 52 and its metabolites in target tissues following intraperitoneal exposure of female rats to doses ranging from 1 to 100 mg/kg BW PCB 52. Three weeks after PCB 52 dosing, PCB 52 remained in adipose, brain, liver, and serum. Several PCB 52 metabolite mixtures were detected in the liver and serum of animals from the 100 mg/kg BW dose groups, with some metabolites only being detected in a single animal. Based on these results, female rats can metabolize PCB 52 to complex PCB metabolite mixtures and likely eliminate the metabolites quickly. The study also found that PCB 52 had minimal effects on certain measurements, such as body and organ weights, serum metabolites, and brain dopamine levels. These effects may be due to the fast elimination of PCB 52. Because PCB 52 is a major contaminant in indoor air, additional work will be needed to determine how age, sex, time of exposure, and exposure route, particularly inhalation, affect PCB 52 disposition and toxicity. Further, it will be important to determine the long-term effects of PCB52 exposure and how this may act synergistically with other factors, such as exposure to other toxicants and diet, to cause disease.

## Funding sources

The National Institute of Environmental Health Sciences/National Institutes of Health supported the project through grants R01ES014901, R01ES031098, P30ES005605, and P42ES013661. The content is solely the responsibility of the authors and does not necessarily represent the official views of the funding agencies listed above.

## Abbreviations

IP: intraperitoneal
PCB: polychlorinated biphenyl
BW: body weight
Nt-LCMS: Nontarget; liquid chromatography-high resolution mass spectrometric
GC-MS/MS: Gas chromatograph mass spectrometric
UPLC-QTof-MS: Ultra-performance liquid chromatography-quadrupole time-of-flight mass spectrometric
QC: quality control
PCA: principle component analysis
ANOVA: analysis of variance
DA: dopamine
DOPAC: 3,4-Dihydroxyphenylacetic acid
DOPAL: 3,4-dihydroxyphenylacetaldehyde
PLS-DA: partial least squares-discriminant analysis (PLS-DA)
VIP: variable importance in projection
CYP: cytochrome P450 enzyme
SD: Sprague-Dawley
MOC: methyoxyamine hydrochloride
TMS: N,O-Bis(trimethylsilyl)trifluoroacetamide
SPE: solid phase extraction
SULT: sulfotransferase
COMT: catechol-O-methyltransferase
ND: non-detect

## Acknowledgments

The authors thank Dr. Andrea Adamcakova-Dodd for helping with the animal study and the Fraternal Order of Eagles Diabetes Research Center Metabolomics Core Facility at the University of Iowa for performing the metabolomic analyses.

## Author contributions

**Bullert:** Data curation; Formal analysis; Investigation; Methodology; Validation; Visualization; Roles/Writing - original draft; Writing - review & editing. **Li:** Data curation; Formal analysis; Investigation; Methodology; Validation; Visualization; Writing - review & editing. **Zhang:** Data curation; Formal analysis; Investigation; Methodology; Validation; Visualization; Writing - review & editing. **Lee:** Formal analysis; Investigation; Validation; Writing - review & editing. **Pulliam:** Conceptualization; Formal analysis; Investigation; Validation; Writing - review & editing. **Cagle:** Formal analysis; Investigation; Validation; Writing - review & editing. **Doorn:** Conceptualization; Funding acquisition; Methodology; Writing - review & editing. **Klingelhutz:** Conceptualization; Investigation; Methodology; Writing - review & editing. **Robertson:** Conceptualization; Funding acquisition; Project administration; Supervision; Writing - review & editing. **Lehmler:** Conceptualization; Funding acquisition; Project administration; Supervision; Validation; Visualization; Roles/Writing - original draft; Writing - review & editing.

## Notes

https://doi.org/10.25820/data.006225

### Competing Interest Statement

The authors have declared no competing interest.

### Summary of Updates

Major changes to the manuscript include adding a Table summarizing the recoveries of the ongoing precision and recovery standard and a description of the measurement of dopamine and its metabolites. The researchers contributing the dopamine data were added as authors to the manuscript. Finally, the language of the manuscript was revised to include references to studies of airborne PCBs from Europe that were not cited in the original version of this manuscript.

## References

Agus, S., Akkaya, H., Daglioglu, N., Eyuboglu, S., Atasayan, O., Mete, F., Colak, C., Sandal, S., Yilmaz, B., 2022. Polychlorinated biphenyls and organochlorine pesticides in breast milk samples and their correlation with dietary and reproductive factors in lactating mothers in Istanbul. Environ Sci Pollut Res Int 29, 3463–3473.

Al Shoyaib, A., Archie, S.R., Karamyan, V.T., 2019. Intraperitoneal route of drug administration: Should it be used in experimental animal studies? Pharm Res 37, 12.

Ampleman, M.D., Martinez, A., DeWall, J., Rawn, D.F., Hornbuckle, K.C., Thorne, P.S., 2015. Inhalation and dietary exposure to PCBs in urban and rural cohorts via congener-specific measurements. Environ Sci Technol 49, 1156–1164.

Anezaki, K., Nakano, T., 2013. Concentration levels and congener profiles of polychlorinated biphenyls, pentachlorobenzene, and hexachlorobenzene in commercial pigments. Environ Sci Pollut Res Int 21, 1–12.

ATSDR, 2000. Toxicological profile for polychlorinated biphenyls.

Azaïs-Braesco, V., Macaire, J.P., Bellenand, P., Robertson, L.W., Pascal, G., 1990. Effects of two prototypic polychlorinated biphenyls (PCBs) on lipid composition of rat liver and serum. J Nutr Biochem 1, 350–354.

Benjamini, Y., Hochberg, Y., 1995. Controlling the false discovery rate: A practical and powerful approach to multiple testing. J R Stat Soc Series B Stat Methodol 57, 289–300.

Birnbaum, L.S., 1994. Endocrine effects of prenatal exposure to PCBs, dioxins, and other xenobiotics: implications for policy and future research. Environ Health Perspect 102, 676–679.

Borlakoglu, J.T., Haegele, K.D., Reich, H.J., Dils, R.R., Wilkins, J.P., 1991. In vitro metabolism of [^14^C]4-chlorobiphenyl and [^14^C]2,2’,5,5’-tetrachlorobiphenyl by hepatic microsomes from rats and pigeons. Evidence against an obligatory arene oxide in aromatic hydroxylation reactions. Int J Biochem 23, 1427–1437.

Borlakoglu, J.T., Wilkins, J.P.G., 1993. Metabolism of di-, tri-, tetra-, penta- and hexachlorobiphenyls by hepatic microsomes isolated from control animals and animals treated with Aroclor 1254, a commercial mixture of polychlorinated biphenyls (PCBs). Comp Biochem Physiol C 105, 95–106.

Bullert, A.J., Doorn, J.A., Stevens, H.E., Lehmler, H.J., 2021. The effects of polychlorinated biphenyl exposure during adolescence on the nervous system: A comprehensive review. Chem Res Toxicol 34, 1948–1952.

Carlson, L.M., Christensen, K., Sagiv, S.K., Rajan, P., Klocke, C.R., Lein, P.J., Coffman, E., Shaffer, R.M., Yost, E.E., Arzuaga, X., Factor-Litvak, P., Sergeev, A., Toborek, M., Bloom, M.S., Trgovcich, J., Jusko, T.A., Robertson, L., Meeker, J., Keating, A.F., Blain, R., Silva, R., Snow, S., Lin, C., Shipkowski, K., Ingle, B., Lehmann, G.M., 2022. A systematic evidence map for the evaluation of noncancer health effects and exposures to polychlorinated biphenyl mixtures. Environ Res, 115148.

Caudle, W.M., Richardson, J.R., Delea, K.C., Guillot, T.S., Wang, M., Pennell, K.D., Miller, G.W., 2006. Polychlorinated biphenyl-induced reduction of dopamine transporter expression as a precursor to Parkinson’s disease-associated dopamine toxicity. Tox Sci 92, 490–499.

Chauhan, K.R., Kodavanti, P.R., McKinney, J.D., 2000. Assessing the role of ortho-substitution on polychlorinated biphenyl binding to transthyretin, a thyroxine transport protein. Tox Appl Pharmacol 162, 10–21.

Chong, J., Soufan, O., Li, C., Caraus, I., Li, S., Bourque, G., Wishart, D.S., Xia, J., 2018. MetaboAnalyst 4.0: towards more transparent and integrative metabolomics analysis. Nucleic Acids Res 46, W486–W494.

Corey, D.A., De Ku, J.L.M., Bingman, V.P., Meserve, L.A., 1996. Effects of exposure to polychlorinated biphenyl (PCB) from conception on growth, and development of endocrine, neurochemical, and cognitive measures in 60 day old rats. Growth Dev Aging 60, 131–143.

De Livera, A.M., Olshansky, G., Simpson, J.A., Creek, D.J., 2018. NormalizeMets: assessing, selecting and implementing statistical methods for normalizing metabolomics data. Metabolomics 14, 54.

Deen, L., Clark, A., Hougaard, K.S., Meyer, H.W., Frederiksen, M., Pedersen, E.B., Petersen, K.U., Flachs, E.M., Bonde, J.P.E., Tottenborg, S.S., 2023. Risk of cardiovascular diseases following residential exposure to airborne polychlorinated biphenyls: A register-based cohort study. Environ Res 222, 115354.

Dhakal, K., Uwimana, E., Adamcakova-Dodd, A., Thorne, P.S., Lehmler, H.J., Robertson, L.W., 2014. Disposition of phenolic and sulfated metabolites after inhalation exposure to 4-chlorobiphenyl (PCB3) in female rats. Chem Res Toxicol 27, 1411–1420.

Doull, J., 2003. The "Red Book" and other risk assessment milestones. Hum Ecol Risk Assess, 1229–1238.

Duffel, M.W., Tuttle, K., Lehmler, H.J., Robertson, L.W., 2021. Human hepatic microsomal sulfatase catalyzes the hydrolysis of polychlorinated biphenyl sulfates: A potential mechanism for retention of hydroxylated PCBs. Environ Toxicol Pharmacol 88, 103757.

Egsmose, E.L., Brauner, E.V., Frederiksen, M., Morck, T.A., Siersma, V.D., Hansen, P.W., Nielsen, F., Grandjean, P., Knudsen, L.E., 2016. Associations between plasma concentrations of PCB 28 and possible indoor exposure sources in Danish school children and mothers. Environ Int 87, 13–19.

Enayah, S.H., Vanle, B.C., Fuortes, L.J., Doorn, J.A., Ludewig, G., 2018. PCB 95 and PCB 153 change dopamine levels and turn-over in PC12 cells. Toxicology 394, 93–101.

Faroon, O., Ruiz, P., 2016. Polychlorinated biphenyls: New evidence from the last decade. Toxicol Ind Health 32, 1825–1847.

Faroon, O.M., Samuel Keith, L., Smith-Simon, C., De Rosa, C.T., 2003. Polychlorinated biphenyls: human health aspects. World Health Organization.

Forgue, T.S., Preston, B.D., Hargraves, W.A., Reich, I.L., Allen, J.R., 1979. Direct evidence that an arene oxide is a metabolic intermediate of 2,2’,5,5’-tetrachlorobiphenyl. Biochem Biophys Res Commun 91, 475–483.

Frame, G.M., 1997. A collaborative study of 209 PCB congeners and 6 Aroclors on 20 different HRGC columns 2. Semi-quantitative Aroclor congener distributions. Anal Bioanal Chem 357, 714–722.

Frame, G.M., Cochran, J.W., Bowadt, S.S., 1996. Complete PCB congener distributions for 17 Aroclor mixtures determined by 3 HRGC systems optimized for comprehensive, quantitative, congener-specific analysis. J High Resolut Chromatogr 19, 657–668.

Gaum, P.M., Esser, A., Schettgen, T., Gube, M., Kraus, T., Lang, J., 2014. Prevalence and incidence rates of mental syndromes after occupational exposure to polychlorinated biphenyls. Int J Hyg Environ Health 217, 765–774.

Gourronc, F.A., Chimenti, M.S., Lehmler, H.J., Ankrum, J.A., Klingelhutz, A.J., 2023. Hydroxylation markedly alters how the polychlorinated biphenyl (PCB) congener, PCB52, affects gene expression in human preadipocytes. Toxicol In Vitro 89, 105568.

Grimm, F.A., He, X.R., Teesch, L.M., Lehmler, H.J., Robertson, L.W., Duffel, M.W., 2015a. Tissue distribution, metabolism, and excretion of 3,3’-dichloro-4’-sulfooxy-biphenyl in the rat. Environ Sci Technol 49, 8087–8095.

Grimm, F.A., Hu, D., Kania-Korwel, I., Lehmler, H.J., Ludewig, G., Hornbuckle, K.C., Duffel, M.W., Bergman, A., Robertson, L.W., 2015b. Metabolism and metabolites of polychlorinated biphenyls. Crit Rev Toxicol 45, 245–272.

Guroff, G., Daly, J.W., Jerina, D.M., Renson, J., Witkop, B., Udenfriend, S., 1967. Hydroxylation-induced migration: the NIH shift. Recent experiments reveal an unexpected and general result of enzymatic hydroxylation of aromatic compounds. Science 157, 1524–1530.

Guvenius, D.M., Hassanzadeh, P., Bergman, A., Noren, K., 2002. Metabolites of polychlorinated biphenyls in human liver and adipose tissue. Environ Toxicol Chem 21, 2264–2269.

Hammel, S.C., Andersen, H.V., Knudsen, L.E., Frederiksen, M., 2023. Inhalation and dermal absorption as dominant pathways of PCB exposure for residents of contaminated apartment buildings. Int J Hyg Environ Health 247, 114056.

Hansen, L.G., Welborn, M.E., 1977. Distribution, dilution and elimination of polychlorinated biphenyl analogs in growing swine. J Pharm Sci 66, 497–501.

Harrad, S., Hazrati, S., Ibarra, C., 2006. Concentrations of polychlorinated biphenyls in indoor air and polybrominated diphenyl ethers in indoor air and dust in Birmingham, United Kingdom: Implications for human exposure. Environ Sci Technol 40, 4633–4638.

Herkert, N.J., Jahnke, J.C., Hornbuckle, K.C., 2018. Emissions of tetrachlorobiphenyls (PCBs 47, 51, and 68) from polymer resin on kitchen cabinets as a non-Aroclor source to residential air. Environ Sci Technol 52, 5154–5160.

Hovander, L., Malmberg, T., Athanasiadou, M., Athanassiadis, I., Rahm, S., Bergman, A., Wehler, E.K., 2002. Identification of hydroxylated PCB metabolites and other phenolic halogenated pollutants in human blood plasma. Arch Environ Contam Toxicol 42, 105–117.

Hu, D., Hornbuckle, K.C., 2010. Inadvertent polychlorinated biphenyls in commercial paint pigments. Environ Sci Technol 44, 2822–2827.

Hu, D., Martinez, A., Hornbuckle, K.C., 2011. Sedimentary records of non-Aroclor and Aroclor PCB mixtures in the Great Lakes. J Great Lakes Res 37, 359–364.

Hutzinger, O., Jamieson, W.D., Safe, S., Paulmann, L., Ammon, R., 1974. Identification of metabolic dechlorination of highly chlorinated biphenyl in rabbit. Nature 252, 698–699.

IACRC, 2016. Polychlorinated biphenyls and polybrominated biphenyls. IARC Monogr Eval Carcinog Risks Hum 107, 9–500.

Jensen, S., Haggberg, L., Jorundsdottir, H., Odham, G., 2003. A quantitative lipid extraction method for residue analysis of fish involving nonhalogenated solvents. J Agric Food Chem 51, 5607–5611.

Jensen, S., Jansson, B., Olsson, M., 1979. Number and identity of anthropogenic substances known to be present in Baltic seals and their possible effects on reproduction. Ann N Y Acad Sci 320, 436–448.

Kaifie, A., Schettgen, T., de Hoogd, M., Kraus, T., Esser, A., 2019. Contamination pathways of polychlorinated biphenyls (PCBs) - From the worker to the family. Int J Hyg Environ Health 222, 1109–1114.

Kania-Korwel, I., Barnhart, C.D., Lein, P.J., Lehmler, H.J., 2015. Effect of pregnancy on the disposition of 2,2’,3,5’,6-pentachlorobiphenyl (PCB95) atropisomers and their hydroxylated metabolites in female mice. Chem Res Toxicol 28, 1774–1783.

Kania-Korwel, I., Barnhart, C.D., Stamou, M., Truong, K.M., El-Komy, M.H., Lein, P.J., Veng-Pedersen, P., Lehmler, H.J., 2012. 2,2’,3,5’,6-Pentachlorobiphenyl (PCB95) and its hydroxylated metabolites are enantiomerically enriched in female mice. Environ Sci Technol 46, 11393–11401.

Kania-Korwel, I., Hornbuckle, K.C., Robertson, L.W., Lehmler, H.J., 2008a. Dose-dependent enantiomeric enrichment of 2,2’,3,3’,6,6’-hexachlorobiphenyl in female mice. Environ Toxicol Chem 27, 299–305.

Kania-Korwel, I., Lukasiewicz, T., Barnhart, C.D., Stamou, M., Chung, H., Kelly, K.M., Bandiera, S., Lein, P.J., Lehmler, H.J., 2017. Editor’s highlight: Congener-specific disposition of chiral polychlorinated biphenyls in lactating mice and their offspring: Implications for PCB developmental neurotoxicity. Tox Sci 158, 101–115.

Kania-Korwel, I., Zhao, H., Norstrom, K., Li, X., Hornbuckle, K.C., Lehmler, H.J., 2008b. Simultaneous extraction and clean-up of polychlorinated biphenyls and their metabolites from small tissue samples using pressurized liquid extraction. J Chromatogr A 1214, 37–46.

Kitamura, S., Jinno, N., Suzuki, T., Sugihara, K., Ohta, S., Kuroki, H., Fujimoto, N., 2005. Thyroid hormone-like and estrogenic activity of hydroxylated PCBs in cell culture. Toxicology 208, 377–387.

Klocke, C., Lein, P.J., 2020. Evidence implicating non-dioxin-like congeners as the key mediators of polychlorinated biphenyl (PCB) developmental neurotoxicity. Int J Mol Sci 21, 1013.

Kofoed, A.B., Deen, L., Hougaard, K.S., Petersen, K.U., Meyer, H.W., Pedersen, E.B., Ebbehoj, N.E., Heitmann, B.L., Bonde, J.P., Tottenborg, S.S., 2021. Maternal exposure to airborne polychlorinated biphenyls (PCBs) and risk of adverse birth outcomes. Eur J Epidemiol 36, 861–872.

Koh, W.X., Hornbuckle, K.C., Marek, R.F., Wang, K., Thorne, P.S., 2016a. Hydroxylated polychlorinated biphenyls in human sera from adolescents and their mothers living in two U.S. Midwestern communities. Chemosphere 147, 389–395.

Koh, W.X., Hornbuckle, K.C., Wang, K., Thorne, P.S., 2016b. Serum polychlorinated biphenyls and their hydroxylated metabolites are associated with demographic and behavioral factors in children and mothers. Environ Int 94, 538–545.

Lam, M.M., Bulow, R., Engwall, M., Giesy, J.P., Larsson, M., 2018. Methylated PACs are more potent than their parent compounds: A study of aryl hydrocarbon receptor-mediated activity, degradability, and mixture interactions in the H4IIE-luc assay. Environ Toxicol Chem 37, 1409–1419.

Lee, D.W., Notter, S.A., Thiruchelvam, M., Dever, D.P., Fitzpatrick, R., Kostyniak, P.J., Cory-Slechta, D.A., Opanashuk, L.A., 2012. Subchronic polychlorinated biphenyl (Aroclor 1254) exposure produces oxidative damage and neuronal death of ventral midbrain dopaminergic systems. Tox Sci 125, 496–508.

Lesmana, R., Shimokawa, N., Takatsuru, Y., Iwasaki, T., Koibuchi, N., 2014. Lactational exposure to hydroxylated polychlorinated biphenyl (OH-PCB 106) causes hyperactivity in male rat pups by aberrant increase in dopamine and its receptor. Environ Toxicol 29, 876–883.

Li, X., Hefti, M.M., Marek, R.F., Hornbuckle, K.C., Wang, K., Lehmler, H.-J., 2022. Assessment of polychlorinated biphenyls and their hydroxylated metabolites in postmortem human brain samples: Age and brain region differences. Environ Sci Technol 56, 9515–9526.

Li, X., Liu, Y., Martin, J.W., Cui, J.Y., Lehmler, H.J., 2021. Nontarget analysis reveals gut microbiome-dependent differences in the fecal PCB metabolite profiles of germ-free and conventional mice. Environ Pollut 268, 115726.

Li, X., Wu, X., Kelly, K.M., Veng-Pedersen, P., Lehmler, H.J., 2019. Toxicokinetics of chiral PCB 136 and its hydroxylated metabolites in mice with a liver-specific deletion of cytochrome P450 reductase. Chem Res Toxicol 32, 727–736.

Li, X., Zhang, C., Wang, K., Lehmler, H.J., 2020. Fatty liver and impaired hepatic metabolism alter the congener-specific distribution of polychlorinated biphenyls (PCBs) in mice with a liver-specific deletion of cytochrome P450 reductase. Environ Pol 266, 115233.

Liberman, D.A., Walker, K.A., Gore, A.C., Bell, M.R., 2020. Sex-specific effects of developmental exposure to polychlorinated biphenyls on neuroimmune and dopaminergic endpoints in adolescent rats. Neurotox Teratol 79, 106880.

Lin, P.H., Sangaiah, R., Ranasinghe, A., Upton, P.B., La, D.K., Gold, A., Swenberg, J.A., 2000. Formation of quinonoid-derived protein adducts in the liver and brain of Sprague-Dawley rats treated with 2,2’,5,5’-tetrachlorobiphenyl. Chem Res Toxicol 13, 710–718.

Liu, Y., Apak, T.I., Lehmler, H.J., Robertson, L.W., Duffel, M.W., 2006. Hydroxylated polychlorinated biphenyls are substrates and inhibitors of human hydroxysteroid sulfotransferase SULT2A1. Chem Res Toxicol 19, 1420–1425.

Liu, Y., Smart, J.T., Song, Y., Lehmler, H.J., Robertson, L.W., Duffel, M.W., 2009. Structure-activity relationships for hydroxylated polychlorinated biphenyls as substrates and inhibitors of rat sulfotransferases and modification of these relationships by changes in thiol status. Drug Metab Dispos 37, 1065–1072.

Lu, Z., Kania-Korwel, I., Lehmler, H.J., Wong, C.S., 2013. Stereoselective formation of mono- and dihydroxylated polychlorinated biphenyls by rat cytochrome P450 2B1. Environ Sci Technol 47, 12184–12192.

Lu, Z., Wong, C.S., 2011. Factors affecting phase I stereoselective biotransformation of chiral polychlorinated biphenyls by rat cytochrome P450 2B1 isozyme. Environ Sci Technol 45, 8298–8305.

Ludewig, G., Lehmann, L., Esch, H., Robertson, L.W., 2008. Metabolic activation of PCBs to carcinogens in vivo - a review. Environ Tox Pharmacol 25, 241–246.

Marek, R.F., Thorne, P.S., Herkert, N.J., Awad, A.M., Hornbuckle, K.C., 2017. Airborne PCBs and OH-PCBs inside and outside urban and rural U.S. schools. Environ Sci Technol 51, 7853–7860.

Mariussen, E., Andersson, P.L., Tysklind, M., Fonnum, F., 2001. Effect of polychlorinated biphenyls on the uptake of dopamine into rat brain synaptic vesicles: a structure-activity study. Toxicol Appl Pharmacol 175, 176–183.

Mariussen, E., Fonnum, F., 2001. The effect of polychlorinated biphenyls on the high affinity uptake of the neurotransmitters, dopamine, serotonin, glutamate and GABA, into rat brain synaptosomes. Toxicology 159, 11–21.

Markowitz, G., Rosner, D., 2018. Monsanto, PCBs, and the creation of a “world-wide ecological problem”. J Public Health Policy 39, 463–540.

Matsusue, K., Ishii, Y., Ariyoshi, N., Oguri, K., 1997. A highly toxic PCB produces unusual changes in the fatty acid composition of rat liver. Toxicol Lett 91, 99–104.

McCann, M.S., Fernandez, H.R., Flowers, S.A., Maguire-Zeiss, K.A., 2021. Polychlorinated biphenyls induce oxidative stress and metabolic responses in astrocytes. Neurotoxicology 86, 59–68.

McLean, M.R., Bauer, U., Amaro, A.R., Robertson, L.W., 1996. Identification of catechol and hydroquinone metabolites of 4-monochlorobiphenyl. Chem Res Toxicol 9, 158–164.

Meerts, I.A., Assink, Y., Cenijn, P.H., Van Den Berg, J.H., Weijers, B.M., Bergman, A., Koeman, J.H., Brouwer, A., 2002. Placental transfer of a hydroxylated polychlorinated biphenyl and effects on fetal and maternal thyroid hormone homeostasis in the rat. Toxicol Sci 68, 361–371.

Mega, O.O., Edesiri, T.P., Victor, E., Kingsley, N.E., Rume, R.A., Faith, F.Y., Simon, O.I., Oghenetega, B.O., Agbonifo-Chijiokwu, E., 2022. d-ribose-l-cysteine abrogates testicular maladaptive responses induced by polychlorinated bisphenol intoxication in rats via activation of the mTOR signaling pathway mediating inhibition of apoptosis, inflammation, and oxidonitrergic flux. J Biochem Mol Toxicol, e23161.

Meyer, H.W., Frederiksen, M., Goen, T., Ebbehoj, N.E., Gunnarsen, L., Brauer, C., Kolarik, B., Muller, J., Jacobsen, P., 2013. Plasma polychlorinated biphenyls in residents of 91 PCB-contaminated and 108 non-contaminated dwellings-an exposure study. Int J Hyg Environ Health 216, 755–762.

Narbonne, J.F., 1979. Effect of polychlorinated biphenyls (Phenoclor DP6) on in vivo protein and RNA synthesis in rat liver. Bull Environ Contam Toxicol 23, 36–43.

Niknam, Y., Feng, W., Cherednichenko, G., Dong, Y., Joshi, S.N., Vyas, S.M., Lehmler, H.J., Pessah, I.N., 2013. Structure-activity relationship of selected meta- and para-hydroxylated non-dioxin like polychlorinated biphenyls: from single RyR1 channels to muscle dysfunction. Toxicol Sci 136, 500–513.

Nomiyama, K., Eguchi, A., Takaguchi, K., Yoo, J., Mizukawa, H., Oshihoi, T., Tanabe, S., Iwata, H., 2019. Targeted metabolome analysis of the dog brain exposed to PCBs suggests inhibition of oxidative phosphorylation by hydroxylated PCBs. Toxicol Appl Pharmacol 377, 114620.

Nomiyama, K., Tsujisawa, Y., Ashida, E., Yachimori, S., Eguchi, A., Iwata, H., Tanabe, S., 2020. Mother to fetus transfer of hydroxylated polychlorinated biphenyl congeners (OH-PCBs) in the Japanese Macaque (*macaca fuscata*): Extrapolation of exposure scenarios to humans. Environ Sci Technol 54, 11386–11395.

Nomiyama, K., Yamamoto, Y., Eguchi, A., Nishikawa, H., Mizukawa, H., Yokoyama, N., Ichii, O., Takiguchi, M., Nakayama, S.M.M., Ikenaka, Y., Ishizuka, M., 2022. Health impact assessment of pet cats caused by organohalogen contaminants by serum metabolomics and thyroid hormone analysis. Sci Total Environ 842, 156490.

Parker, V.S., Squirewell, E.J., Lehmler, H.J., Robertson, L.W., Duffel, M.W., 2018. Hydroxylated and sulfated metabolites of commonly occurring airborne polychlorinated biphenyls inhibit human steroid sulfotransferases SULT1E1 and SULT2A1. Environ Toxicol Pharmacol 58, 196–201.

Parkinson, A., Safe, S.H., Robertson, L.W., Thomas, P.E., Ryan, D.E., Reik, L.M., Levin, W., 1983. Immunochemical quantitation of cytochrome P450 isozymes and epoxide hydrolase in liver microsomes from polychlorinated or polybrominated biphenyl-treated rats. A study of structure-activity relationships. J Biol Chem 258, 5967–5976.

Pessah, I.N., Cherednichenko, G., Lein, P.J., 2010. Minding the calcium store: Ryanodine receptor activation as a convergent mechanism of PCB toxicity. Pharmacol Ther 125, 260–285.

Pierucci, F., Frati, A., Squecco, R., Lenci, E., Vicenti, C., Slavik, J., Francini, F., Machala, M., Meacci, E., 2017. Non-dioxin-like organic toxicant PCB 153 modulates sphingolipid metabolism in liver progenitor cells: its role in Cx43-formed gap junction impairment. Arch Toxicol 91, 749–760.

Quinete, N., Esser, A., Kraus, T., Schettgen, T., 2017. PCB28 metabolites elimination kinetics in human plasma on a real case scenario: Study of hydroxylated polychlorinated biphenyl (OH-PCB) metabolites of PCB28 in a highly exposed German Cohort. Toxicol Lett 276, 100–107.

Quinete, N., Schettgen, T., Bertram, J., Kraus, T., 2014. Occurrence and distribution of PCB metabolites in blood and their potential health effects in humans: A review. Environ Sci Pollut Res Int 21, 11951–11972.

Rodriguez, E.A., Vanle, B.C., Doorn, J.A., Lehmler, H.J., Robertson, L.W., Duffel, M.W., 2018. Hydroxylated and sulfated metabolites of commonly observed airborne polychlorinated biphenyls display selective uptake and toxicity in N27, SH-SY5Y, and HepG2 cells. Environ Toxicol Pharmacol 62, 69–78.

Safe, S., Hutzinger, O., Jones, D., 1975. The mechanism of chlorobiphenyl metabolism. J Agric Food Chem 23, 851–853.

Saktrakulkla, P., Lan, T., Hua, J., Marek, R.F., Thorne, P.S., Hornbuckle, K.C., 2020. Polychlorinated biphenyls in food. Environ Sci Technol 54, 11443–11452.

Schecter, A., Colacino, J., Haffner, D., Patel, K., Opel, M., Papke, O., Birnbaum, L., 2010. Perfluorinated compounds, polychlorinated biphenyls, and organochlorine pesticide contamination in composite food samples from Dallas, Texas, USA. Environ Health Perspect 118, 796–802.

Seegal, R.F., Bush, B., Brosch, K.O., 1994. Decreases in dopamine concentrations in adult, non-human primate brain persist following removal from polychlorinated biphenyls. Toxicology 86, 71–87.

Seegal, R.F., Bush, B., Shain, W., 1990. Lightly chlorinated ortho-substituted PCB congeners decrease dopamine in nonhuman primate brain and in tissue culture. Toxicol Appl Pharmacol 106, 136–144.

Seelbach, M., Chen, L., Powell, A., Choi, Y.J., Zhang, B., Hennig, B., Toborek, M., 2010. Polychlorinated biphenyls disrupt blood-brain barrier integrity and promote brain metastasis formation. Environ Health Perspect 118, 479–484.

Sethi, S., Morgan, R.K., Feng, W., Lin, Y., Li, X., Luna, C., Koch, M., Bansal, R., Duffel, M.W., Puschner, B., Zoeller, R.T., Lehmler, H.J., Pessah, I.N., Lein, P.J., 2019. Comparative analyses of the 12 most abundant PCB congeners detected in human maternal serum for activity at the thyroid hormone receptor and ryanodine receptor. Environ Sci Technol 53, 3948–3958.

Shimada, T., Kakimoto, K., Takenaka, S., Koga, N., Uehara, S., Murayama, N., Yamazaki, H., Kim, D., Guengerich, F.P., Komori, M., 2016. Roles of human CYP2A6 and monkey CYP2A24 and 2A26 cytochrome P450 enzymes in the oxidation of 2,5,2’,5’-tetrachlorobiphenyl. Drug Metab Dispos 44, 1899–1909.

Song, Y., Wagner, B.A., Lehmler, H.J., Buettner, G.R., 2008. Semiquinone radicals from oxygenated polychlorinated biphenyls: Electron paramagnetic resonance studies. Chem Res Toxicol 21, 1359–1367.

Stamou, M., Uwimana, E., Flannery, B.M., Kania-Korwel, I., Lehmler, H.J., Lein, P.J., 2015. Subacute nicotine co-exposure has no effect on 2,2’,3,5’,6-pentachlorobiphenyl disposition but alters hepatic cytochrome P450 expression in the male rat. J Toxicol 338, 59–68.

Stockholm-Convention, 2019. Stockholm convention overview of PCBs. http://chm.pops.int/Implementation/IndustrialPOPs/PCBs/Overview/tabid/273/Default.aspx.

Sundström, G., Jansson, B., 1975. The metabolism of 2,2’,3,5’,6-pentachlorobiphenyl in rats, mice and quails. Chemosphere 4, 361–370.

Tan, Y., Chen, C.H., Lawrence, D., Carpenter, D.O., 2004. Ortho-substituted PCBs kill cells by altering membrane structure. Toxicol Sci 80, 54–59.

Tehrani, R., Van Aken, B., 2014. Hydroxylated polychlorinated biphenyls in the environment: sources, fate, and toxicities. Environ Sci Pollut Res Int 21, 6334–6345.

Tompkins, S.C., Sheldon, R.D., Rauckhorst, A.J., Noterman, M.F., Solst, S.R., Buchanan, J.L., Mapuskar, K.A., Pewa, A.D., Gray, L.R., Oonthonpan, L., Sharma, A., Scerbo, D.A., Dupuy, A.J., Spitz, D.R., Taylor, E.B., 2019. Disrupting mitochondrial pyruvate uptake directs glutamine into the TCA cycle away from glutathione synthesis and impairs hepatocellular tumorigenesis. Cell Rep 28, 2608–2619 e2606.

Totland, C., Nerdal, W., Steinkopf, S., 2016. Effects and location of coplanar and noncoplanar PCB in a lipid bilayer: A solid-state NMR study. Environ Sci Technol 50, 8290–8295.

Uwimana, E., Cagle, B., Yeung, C., Li, X., Patterson, E.V., Doorn, J.A., Lehmler, H.J., 2019. Atropselective oxidation of 2,2’,3,3’,4,6’-hexachlorobiphenyl (PCB 132) to hydroxylated metabolites by human liver microsomes and its implications for PCB 132 neurotoxicity. Toxicol Sci 171, 406–420.

Uwimana, E., Li, X., Lehmler, H.J., 2018. Human liver microsomes atropselectively metabolize 2,2’,3,4’,6-pentachlorobiphenyl (PCB 91) to a 1,2-shift product as the major metabolite. Environ Sci Technol 52, 6000–6008.

Wang, H., Adamcakova-Dodd, A., Lehmler, H.J., Hornbuckle, K.C., Thorne, P.S., 2022. Toxicity assessment of 91-day repeated inhalation exposure to an indoor school air mixture of PCBs. Environ Sci Technol 56, 1780–1790.

Wu, X., Barnhart, C., Lein, P.J., Lehmler, H.J., 2015. Hepatic metabolism affects the atropselective disposition of 2,2’,3,3’,6,6’-hexachlorobiphenyl (PCB136) in mice. Environ Sci Technol 49, 616–625.

Wu, X., Zhai, G., Schnoor, J.L., Lehmler, H.J., 2020. Atropselective disposition of 2,2’,3,4’,6-pentachlorobiphenyl (PCB91) and identification of its metabolites in mice with liver-specific deletion of cytochrome P450 reductase. Chem Res Toxicol 33, 1328–1338.

Yoshida, S., Nakamura, A., 1977. Studies on a metabolite of PCB (methylsulfone PCB) in mother milk. J Food Hyg Saf 18, 387–388.

Zhang, C.-Y., Flor, S., Ruiz, P., Dhakal, R., Hu, X., Teesch, L.M., Ludewig, G., Lehmler, H.-J., 2020. 3,3’-dichlorobiphenyl is metabolized to a complex mixture of oxidative metabolites, including novel methoxylated metabolites, by HepG2 cells. Environ Sci Technol 54, 12345–12357.

Zhang, C.Y., Klocke, C.R., Lein, P.J., Lehmler, H.J., 2021. Disposition of PCB 11 in mice following acute oral exposure. Chem Res Toxicol 34, 988–991.

Zhang, C.Y., Li, X., Flor, S., Ruiz, P., Kruve, A., Ludewig, G., Lehmler, H.J., 2022a. Metabolism of 3-chlorobiphenyl (PCB2) in a human-relevant cell line: Evidence of dechlorinated metabolites. Environ Sci Technol 56, 12460–12472.

Zhang, D., Saktrakulkla, P., Marek, R.F., Lehmler, H.J., Wang, K., Thorne, P.S., Hornbuckle, K.C., Duffel, M.W., 2022b. PCB sulfates in serum from mothers and children in urban and rural U.S. communities. Environ Sci Technol 56, 6537–6547.

