## Supplementary Material for "Disposition and Metabolomic Effects of 2,2’,5,5’-Tetrachlorobiphenyl in Female Rats Following Intraperitoneal Exposure"

^a^ Department of Occupational and Environmental Health, College of Public Health, University of Iowa, Iowa City, IA 52242, USA; ^b^ Interdisciplinary Graduate Program in Neuroscience, University of Iowa, Iowa City, IA 52242, USA; ^c^ Interdisciplinary Program in Human Toxicology, University of Iowa, Iowa City, IA 52242, USA; ^d^ Department of Pharmaceutical Sciences and Experimental Therapeutics, University of Iowa, Iowa City, IA 52242, USA; ^e^ Department of Microbiology and Immunology, Carver College of Medicine, University of Iowa, Iowa City, IA 52242, USA

Corresponding Author:

Dr. Hans-Joachim Lehmler

The University of Iowa

Department of Occupational and Environmental Health

University of Iowa Research Park, #221 IREH

Iowa City, IA 52242-5000

**Table of Contents**

| Chemicals | S4 |
| --- | --- |
| **Table S1.** Abbreviations and unique identifiers of the test compounds and all analytical standards used in this study. | S5 |
| **Table S2.** GC/MS/MS information regarding internal standards, surrogate standards, and quantitative standards. Included is the precursor and product ions along with the corresponding collision energy for each analyte included. | S6 |
| **Table S3.** Percent recoveries of surrogate standards for each tissue. | S7 |
| **Table S4.** Ongoing precision recovery recoveries (%) of PCB 52 and 4-OH-PCB 52 for each tissue. | S8 |
| **Table S5.** Method detection limit (ng) and limit of detection for each tissue type (ng/g) for gas chromatographic quantifications of PCB 52 and 4-OH-PCB 52. | S9 |
| **Table S6.** Tissue weights for each exposure group. Data are presented as the mean (SD). | S10 |
| **Table S7.** Levels (ng/g) of PCB 52 or the major metabolite, 4-OH-PCB 52, in each tissue for each PCB 52 exposure group. Values are represented as the mean (SD). | S11 |
| **Fig. S1.** Body weight-adjusted spleen weights. | S12 |
| **Fig. S2**. Body weight-adjusted thymus weights. | S13 |
| **Fig. S3.** One tetrachlorinated OH-PCB metabolite was detected by LC-Orbitrap MS in the liver of PCB 52 exposed rats. | S14 |
| **Fig. S4.** Two tetrachlorinated PCB sulfates were detected by LC-Orbitrap MS in the liver of PCB 52 exposed rats. | S15 |
| **Fig. S5.** One tetrachlorinated diOH-PCB was detected by LC-Orbitrap MS in the liver of PCB 52 exposed rats. | S16 |
| **Fig. S6.** Three tetrachlorinated OH-PCB sulfates were detected by LC-Orbitrap MS in the liver of PCB 52 exposed rats. | S17 |
| **Fig. S7.** One tetrachlorinated MeO-OH-PCB was detected by LC-Orbitrap MS in the liver of PCB 52 exposed rats. | S18 |
| **Fig. S8.** Four tetrachlorinated MeO-OH-PCB sulfates were detected by LC-Orbitrap MS in the liver of PCB 52 exposed rats. | S19 |
| **Fig. S9.** One trichlorinated OH-PCB sulfate was detected by LC-Orbitrap MS in the liver of PCB 52 exposed rats. | S20 |
| **Fig. S10.** One trichlorinated MeO-PCB sulfate was detected by LC-Orbitrap MS in the liver of PCB 52 exposed rats. | S21 |
| **Fig. S11.** One trichlorinated MeO-diOH-PCB was tentatively detected by LC-Orbitrap MS in the liver of PCB 52 exposed rats. | S22 |
| **Fig. S12.** One trichlorinated MeO-OH-PCB sulfate was detected by LC-Orbitrap MS in the liver of PCB 52 exposed rats. | S23 |
| **Fig. S13.** One tetrachlorinated OH-PCB was detected by LC-Orbitrap MS in the serum of PCB 52 exposed rats. | S24 |
| **Fig. S14.** One tetrachlorinated PCB sulfate was detected by LC-Orbitrap MS in the serum of PCB 52 exposed rats. | S25 |
| **Fig. S15.** Two tetrachlorinated OH-PCB sulfates were detected by LC-Orbitrap MS in the serum of PCB 52 exposed rats. | S26 |
| **Fig. S16.** One tetrachlorinated MeO-OH-PCB was detected by LC-Orbitrap MS in the serum of PCB 52 exposed rats. | S27 |
| **Fig. S17.** One tetrachlorinated MeO-OH-PCB sulfate was tentatively detected by LC-Orbitrap MS in the serum of PCB 52 exposed rats. | S28 |
| **Fig. S18.** Two trichlorinated MeO-OH-PCB sulfates were tentatively detected by LC-Orbitrap MS in the serum of PCB 52 exposed rats. | S29 |

**Chemicals.** The test compound, PCB 52, was synthesized and authenticated using a published guideline (Li et al., 2018). Details regarding its authentication are reported elsewhere (Saktrakulkla et al., 2022; Sethi et al., 2019). Analytical standards for gas chromatographic analyses, including 3,3',4,4'-tetrachlorobiphenyl (PCB 77) and 2,5,2',5'-tetrachlorobiphenyl-4-ol (4-OH-PCB 52) were synthesized and authenticated as described previously (Rodriguez et al., 2016; Sethi et al., 2019; Shaikh et al., 2008; Tampal et al., 2002). 2,3,3',4,5,5'-Hexachlorobiphenyl-4'-ol (4'-OH-PCB 159) and 2,2',3,4,4',5,6,6'-octachlorobiphenyl (PCB 204) were purchased from AccuStandard, Inc (New Haven, Connecticut, USA). PCB 77 and 4'-OH-PCB 159 were added to all samples as surrogate recovery standards. PCB 204 was used as an internal standard to adjust for volume differences between samples. The recovery standard for the Nt-LCMS analysis, 4'-chloro-3'-fluoro-4-hydroxy-biphenyl (F-OH-PCB 3) was prepared and authenticated as reported elsewhere (Dhakal et al., 2012; Li et al., 2022). The potassium salt of perfluorooctanesulfonic acid, the internal standard for the Nt-LCMS analysis, was provided by Fisher Scientific (Pittsburg, Pennsylvania, USA).

**Table S1.** Abbreviations and unique identifiers of the test compounds and all analytical standards used in this study.

| **Abbreviation** | **IUPAC Name** | **FORMULA** | **Isomeric SMILES** | **InChI** | **InChIKey** | **CAS Registry Number** | **CAS Registry URL** | **PubChem CID** | **PubChem**  **Link** | **DTXSID** | **Comptox**  **Link** |
| --- | --- | --- | --- | --- | --- | --- | --- | --- | --- | --- | --- |
| 3-F-4'-OH-PCB 3 | 4-(4-chloro-3-fluorophenyl)phenol | C12H8ClFO | C1=CC(=CC=C1C2=CC(=C(C=C2)Cl)F)O | InChI=1S/C12H8ClFO/c13-11-6-3-9(7-12(11)14)8-1-4-10(15)5-2-8/h1-7,15H | DBNVNCYZHMXJJL-UHFFFAOYSA-N | 893736-99-7 | https://commonchemistry.cas.org/detail?cas_rn=893736-99-7 | 20099942 | https://pubchem.ncbi.nlm.nih.gov/compound/20099942 | NA | NA |
| PCB 30 | 2,4,6-Trichlorobiphenyl | C12H7Cl3 | C1=CC=C(C=C1)C2=C(C=C(C=C2Cl)Cl)Cl | InChI=1S/C12H7Cl3/c13-9-6-10(14)12(11(15)7-9)8-4-2-1-3-5-8/h1-7H | MTLMVEWEYZFYTH-UHFFFAOYSA-N | 35693-92-6 | https://commonchemistry.cas.org/detail?cas_rn=35693-92-6 | 37247 | https://pubchem.ncbi.nlm.nih.gov/compound/37247 | DTXSID7073482 | https://comptox.epa.gov/dashboard/chemical/details/DTXSID7073482 |
| PCB 52 | 2,2',5,5'-Tetrachlorobiphenyl | C12H6Cl4 | C1=CC(=C(C=C1Cl)C2=C(C=CC(=C2)Cl)Cl)Cl | InChI=1S/C12H6Cl4/c13-7-1-3-11(15)9(5-7)10-6-8(14)2-4-12(10)16/h1-6H | HCWZEPKLWVAEOV-UHFFFAOYSA-N | 35693-99-3 | https://commonchemistry.cas.org/detail?cas_rn=35693-99-3 | 37248 | https://pubchem.ncbi.nlm.nih.gov/compound/37248 | DTXSID3038305 | https://comptox.epa.gov/dashboard/chemical/details/DTXSID3038305 |
| 4-OH-PCB 52 | 2,5-dichloro-4-(2,5-dichlorophenyl)phenol | C12H6Cl4O | C1=CC(=C(C=C1Cl)C2=CC(=C(C=C2Cl)O)Cl)Cl | InChI=1S/C12H6Cl4O/c13-6-1-2-9(14)7(3-6)8-4-11(16)12(17)5-10(8)15/h1-5,17H | ZKDSNFDCQYBBIU-UHFFFAOYSA-N | 51274-68-1 | https://commonchemistry.cas.org/detail?cas_rn=51274-68-1 | 39971 | https://pubchem.ncbi.nlm.nih.gov/compound/39971 | DTXSID10199272 | https://comptox.epa.gov/dashboard/DTXSID10199272 |
| PCB 77 | 3,3',4,4'-Tetrachlorobiphenyl | C12H6Cl4 | C1=CC(=C(C=C1C2=CC(=C(C=C2)Cl)Cl)Cl)Cl | InChI=1S/C12H6Cl4/c13-9-3-1-7(5-11(9)15)8-2-4-10(14)12(16)6-8/h1-6H | UQMGJOKDKOLIDP-UHFFFAOYSA-N | 32598-13-3 | https://commonchemistry.cas.org/detail?cas_rn=32598-13-3 | 36187 | https://pubchem.ncbi.nlm.nih.gov/compound/36187 | DTXSID5022514 | https://comptox.epa.gov/dashboard/chemical/details/DTXSID5022514 |
| 4'-OH-PCB 79 | 2,6-dichloro-4-(3,4-dichlorophenyl)phenol | C12H6Cl4O | C1=CC(=C(C=C1C2=CC(=C(C(=C2)Cl)O)Cl)Cl)Cl | InChI=1S/C12H6Cl4O/c13-8-2-1-6(3-9(8)14)7-4-10(15)12(17)11(16)5-7/h1-5,17H | RQGVZEFZWFEKQR-UHFFFAOYSA-N | 111810-41-4 | https://commonchemistry.cas.org/detail?cas_rn=111810-41-4 | 119346 | https://pubchem.ncbi.nlm.nih.gov/compound/119346 | DTXSID60149742 | https://comptox.epa.gov/dashboard/DTXSID60149742 |
| 4'-OH-PCB 159 | 2,6-dichloro-4-(2,3,4,5-tetrachlorophenyl)phenol | C12H4Cl6O | C1=C(C=C(C(=C1Cl)O)Cl)C2=CC(=C(C(=C2Cl)Cl)Cl)Cl | InChI=1S/C12H4Cl6O/c13-6-3-5(9(16)11(18)10(6)17)4-1-7(14)12(19)8(15)2-4/h1-3,19H | PZAKBNHYWBSZAF-UHFFFAOYSA-N | 158076-63-2 | https://chem.nlm.nih.gov/chemidplus/sid/0158076632 | 178005 | https://pubchem.ncbi.nlm.nih.gov/compound/178005 | DTXSID70166369 | https://comptox.epa.gov/dashboard/DTXSID70166369 |
| PCB 204 | 2,2',3,4,4',5,6,6'-Octachlorobiphenyl | C12H2Cl8 | C1=C(C=C(C(=C1Cl)C2=C(C(=C(C(=C2Cl)Cl)Cl)Cl)Cl)Cl)Cl | InChI=1S/C12H2Cl8/c13-3-1-4(14)6(5(15)2-3)7-8(16)10(18)12(20)11(19)9(7)17/h1-2H | JDZUWXRNKHXZFE-UHFFFAOYSA-N | 74472-52-9 | https://commonchemistry.cas.org/detail?cas_rn=74472-52-9 | 91721 | https://pubchem.ncbi.nlm.nih.gov/compound/91721 | DTXSID7074240 | https://comptox.epa.gov/dashboard/chemical/details/DTXSID7074240 |

**Table S2.** GC/MS/MS information regarding internal standards, surrogate standards, and quantitative standards. Included is the precursor and product ions along with the corresponding collision energy for each analyte included.

| **Analyte** | **Precursor Ion (*m/z*)** | **Product Ion (*m/z*)** | **Collision Energy (eV)** |
| --- | --- | --- | --- |
| PCB 52 | 291.9 | 222.0 | 25 |
| 4-OH-PCB 52 | 321.9 | 278.9 | 20 |
| PCB 77 | 291.9 | 222.0 | 25 |
| PCB 204 | 429.7 | 357.8 | 35 |
| 4'-OH-PCB 159 | 389.9 | 374.9 | 15 |

**Table S3.** Percent recoveries of surrogate standards for each tissue.

| **Recovery Standard** | **Adipose**, N = 30 | **Brain**, N = 34 | **Liver**, N = 33 | **Serum**, N = 33 |
| --- | --- | --- | --- | --- |
| PCB 77 |  |  |  |  |
| Mean (SD) | 82 (12) | 96 (17) | 85 (18) | 99 (30) |
| Range | 51, 105 | 70, 141 | 59, 127 | 59, 165 |
| 4'-OH-PCB 159 |  |  |  |  |
| Mean (SD) | 83 (16) | 108 (21) | 89 (14) | 87 (10) |
| Range | 63, 131 | 61, 155 | 61, 123 | 67, 110 |

N represents the number of samples analyzed. Samples included from three pools of samples analyzed concurrently. PCB 77, a tetra chlorinated biphenyl, was used as a surrogate standard for PCB 52. 4'-OH-PCB 159 was used as a surrogate standard for hydroxylated PCB metabolites to determine extraction efficiency.

**Table S4.** Ongoing precision recovery recoveries (%) of PCB 52 and 4-OH-PCB 52 for each tissue.

| **PCB or Metabolite** | **Blank Spike** | | | | | | | **Tissue Blank Spike** | | | | | | |
| --- | --- | --- | --- | --- | --- | --- | --- | --- | --- | --- | --- | --- | --- | --- |
|  | **Adipose**,  N = 4 | **Brain**,  N = 6 |  |  | **Liver**,  N = 9 |  | **Serum**,  N = 6 | **Adipose**,  N = 4 | **Brain**,  N = 6 |  |  | **Liver**,  N = 9 |  | **Serum**,  N = 6 |
| PCB 52 | 85 (27) | 99 (18) |  |  | 86 (27) |  | 80 (21) | 98 (31) | 94 (13) |  |  | 92 (30) |  | 85 (23) |
| 4-OH-PCB 52 | 78 (11) | 83 (12) |  |  | 77 (9) |  | 83 (22) | 86 (16) | 75 (12) |  |  | 87 (12) |  | 75 (17) |

N represents the number of samples analyzed.

**Table S5.** Method detection limit (ng) and limit of detection for each tissue type (ng/g) for gas chromatographic quantifications of PCB 52 and 4-OH-PCB 52.

| **Tissue LODs** | **PCB 52** | **4-OH-PCB 52** | **X1-PCB 52** |
| --- | --- | --- | --- |
| **MDL [ng]^1^** | 0.08 | 0.07 | 0.04 |
| **Adipose** | | | |
| LOD^2^ [ng] | 0.09 | 0.03 | 0.02 |
| LOD^3^ [ng/g] | 0.9 | 0.34 | 0.15 |
| **Brain** | | | |
| LOD^2^ [ng] | 0.12 | 0.1 | 0.14 |
| LOD^3^ [ng/g] | 0.17 | 0.16 | 0.24 |
| **Liver** | | | |
| LOD^2^ [ng] | 0.02 | 0.01 | 0.01 |
| LOD^3^ [ng/g] | 0.04 | 0.03 | 0.02 |
| **Serum** | | | |
| LOD^2^ [ng] | 0.08 | 0.04 | 0.03 |
| LOD^3^ [ng/g] | 0.28 | 0.09 | 0.06 |

^1^ MDL, method detection limit, was calculated based on method blanks.

^2^ LOD, limit of detection, was calculated based on matrix blanks for each tissue using the formula LOD= mean_blank_ + t_0.01,n= 8_ *SD_blank_, where mean_blank_ is the mean of eight blank measures, t_0.01,n= 8_ is Student’s t-value for n – 1 degrees of freedom at the 99% confidence level, and SD_blank_ is the standard deviation of the blank measures.

^3^ LOD adjusted for tissue weight (g).

**Table S6.** Tissue weights for each exposure group. Data are presented as the mean (SD).

| **Tissue** | **0 mg/kg BW,**  **N = 3** | **1 mg/kg BW,**  **N = 3** | **10 mg/kg BW,**  **N = 3** | **100 mg/kg BW,**  **N = 5** |
| --- | --- | --- | --- | --- |
| Body weight (g) | 179 (9) | 176 (6) | 173 (11) | 175 (12) |
| Spleen (g) | 0.50 (0.02) | 0.43 (0.05) | 0.50 (0.07) | 0.47 (0.06) |
| Liver (g) | 7.86 (0.57) | 7.31 (0.25) | 6.78 (0.29) | 7.22 (0.65) |
| Thymus (g) | 0.52 (0.07) | 0.48 (0.02) | 0.48 (0.01) | 0.45 (0.05) |
| Body weight adjusted spleen (g/g bw) | 0.0028 (0.0001) | 0.0025 (0.0003) | 0.0029 (0.0002) | 0.0027 (0.0002) |
| Body weight adjusted liver (g/g bw) | 0.0438 (0.0013) | 0.0416 (0.0003) | 0.0392 (0.0010) | 0.0417 (0.0009) |
| Body weight adjusted thymus (g/g bw) | 0.0029 (0.0003) | 0.0027 (0.0001) | 0.0028 (0.0002) | 0.0026 (0.0003) |
| SD, standard deviation, bw, body weight. | | | | |

**Table S7.** Levels (ng/g) of PCB 52 or the major metabolite, 4-OH-PCB 52, in each tissue for each PCB 52 exposure group. Values are represented as the mean (SD).

| **Analyte** | **Vehicle** | | | | **1 mg/kg BW** | | | |
| --- | --- | --- | --- | --- | --- | --- | --- | --- |
|  | **Adipose,**  **N = 3** | **Brain,**  **N = 3** | **Liver,**  **N = 3** | **Serum,**  **N = 3** | **Adipose,**  **N = 3** | **Brain,**  **N = 3** | **Liver,**  **N = 3** | **Serum,**  **N = 3** |
| PCB 52 | NA (NA) | NA (NA) | NA (NA) | NA (NA) | 55 (51) | 1 (1) | 1 (1) | 1 (1) |
| 4-OH-PCB 52 | NA (NA) | NA (NA) | NA (NA) | NA (NA) | NA (NA) | NA (NA) | NA (NA) | 0.140 (0.042) |
| X1-PCB 52 | NA (NA) | NA (NA) | NA (NA) | NA (NA) | NA (NA) | NA (NA) | NA (NA) | NA (NA) |

| **Analyte** |  | | | |  | | | | **10 mg/kg BW** | | | | **100 mg/kg BW** | | | |
| --- | --- | --- | --- | --- | --- | --- | --- | --- | --- | --- | --- | --- | --- | --- | --- | --- |
|  |  |  |  |  |  |  |  |  | **Adipose,**  **N = 3** | **Brain,**  **N = 3** | **Liver,**  **N = 3** | **Serum,**  **N = 3** | **Adipose,**  **N = 5** | **Brain,**  **N = 5** | **Liver,**  **N = 5** | **Serum,**  **N = 5** |
| PCB 52 |  |  |  |  |  |  |  |  | 357 (186) | 7 (5) | 6 (4) | 4 (3) | 2,018 (3,255) | 20 (35) | 28 (37) | 15 (26) |
| 4-OH-PCB 52 |  |  |  |  |  |  |  |  | NA (NA) | NA (NA) | 0.15 (0.06) | 0.70 (0.39) | 1.60 (NA) | 0.17 (NA) | 0.57 (0.75) | 2.27 (3.72) |
| X1-PCB 52 |  |  |  |  |  |  |  |  | NA (NA) | NA (NA) | 0.28 (0.02) | 0.07 (NA) | 1.24 (1.87) | 0.24 (NA) | 0.85 (0.95) | 0.69 (1.04) |

SD, standard deviation; NA, not available.X1, unknown monohydroxylated compound; BW, Body weight,

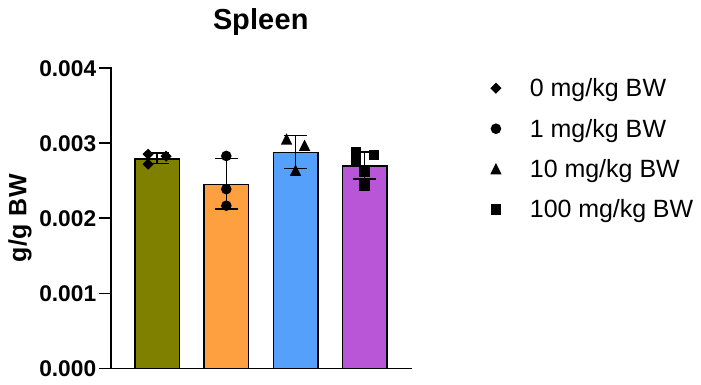

**Figure S1.** Body weight-adjusted spleen weights. Spleen weights were not significantly affected after 3 weeks post IP injection of PCB 52. Group means were compared using one-way ANOVA followed by Tukey’s pairwise comparisons.

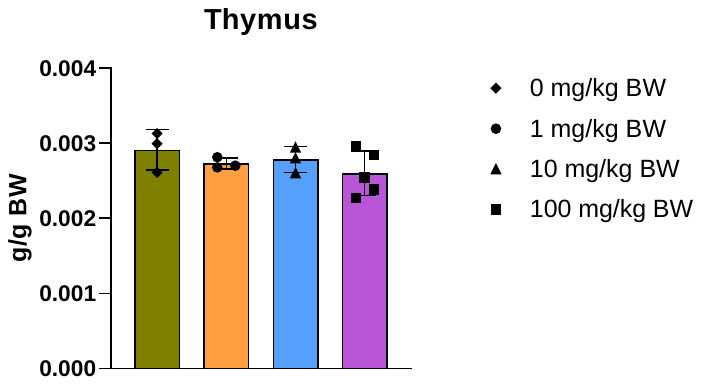

**Figure S2.** Body weight-adjusted thymus weights. Thymus weights were not significantly different 3 weeks post IP injection of PCB 52. Group means were compared using one-way ANOVA followed by Tukey’s pairwise comparisons.

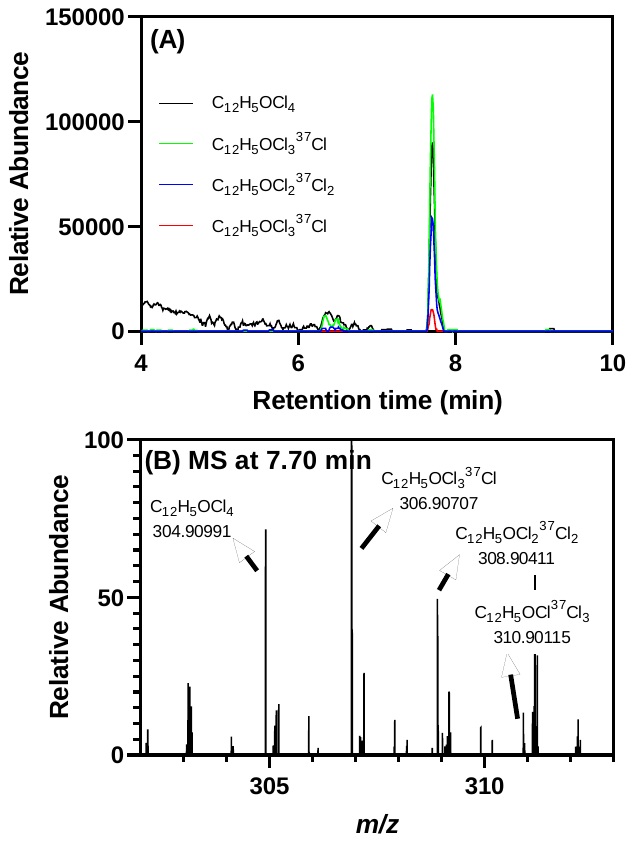

**Figure S3.** One tetrachlorinated OH-PCB metabolite was detected by LC-Orbitrap MS in the liver of PCB 52 exposed rats. (A) Chromatograms extracted based on the theoretical accurate mass of the top four high-abundance isotope ions of tetrachlorinated OH-PCBs ([C_12_H_5_OCl_4_]^-^, *m/z* 304.91000 for the monoisotopic ion) show a peak at 7.70 min. (B) The accurate masses of four high-abundance isotope ions at 7.70 min match the theoretical accurate mass and isotopic pattern of a tetrachlorinated compound. The LC-Orbitrap MS analysis was performed in the negative polarity mode.

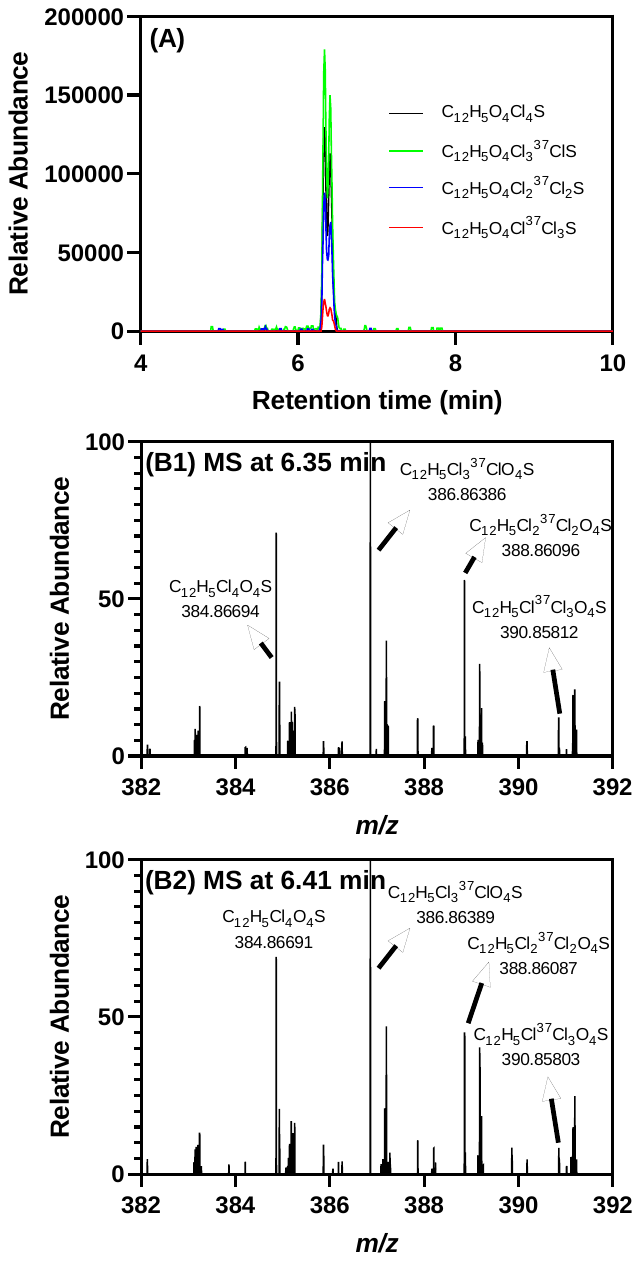

**Figure S4.** Two tetrachlorinated PCB sulfates were detected by LC-Orbitrap MS in the liver of PCB 52 exposed rats. (A) Chromatograms extracted based on the theoretical accurate mass of the top four high-abundance isotope ions of tetrachlorinated PCB sulfates ([C_12_H_5_Cl_4_O_4_S]^-^, *m/z* 384.86681 for the monoisotopic ion) show peaks at 6.35 and 6.41 min. The accurate masses of four high-abundance isotope ions at (B1) 6.35 min and (B2) 6.41 min match the theoretical accurate mass and isotopic pattern of a tetrachlorinated compound. The LC-Orbitrap MS analysis was performed in the negative polarity mode.

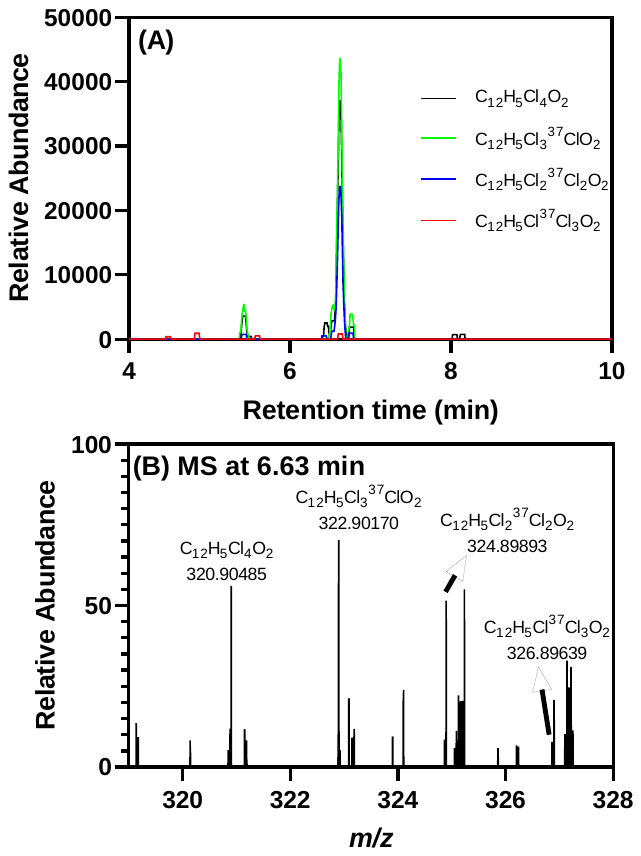

**Figure S5.** One tetrachlorinated diOH-PCB was detected by LC-Orbitrap MS in the liver of PCB 52 exposed rats. (A) Chromatograms extracted based on the theoretical accurate mass of the top four high-abundance isotope ions of tetrachlorinated diOH-PCBs ([C_12_H_5_Cl_4_O_2_]^-^, *m/z* 320.90491 for the monoisotopic ion) show a peak at 6.63 min. (B) The accurate masses of four high-abundance isotope ions at 6.63 min match the theoretical accurate mass and isotopic pattern of a tetrachlorinated compound. The LC-Orbitrap MS analysis was performed in the negative polarity mode.

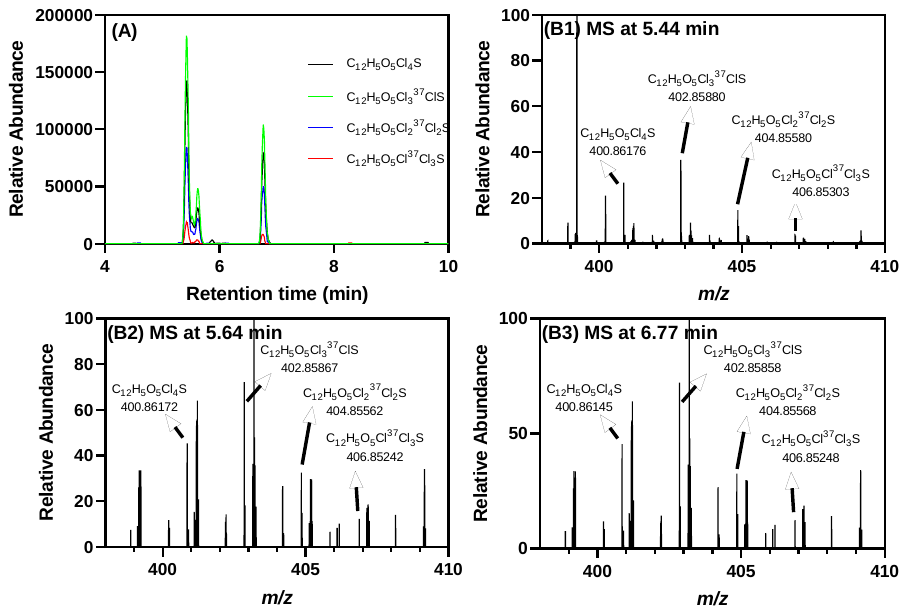

**Figure S6.** Three tetrachlorinated OH-PCB sulfates were detected by LC-Orbitrap MS in the liver of PCB 52 exposed rats. (A) Chromatograms extracted based on the theoretical accurate mass of the top four high-abundance isotope ions of tetrachlorinated OH-PCB sulfates ([C_12_H_5_O_5_Cl_4_S]^-^, *m/z* 400.86173 for the monoisotopic ion) show peaks at 5.44, 5.64, and 6.77 min. The accurate masses of several high-abundance isotope ions at (B1) 5.44 min, (B2) 5.64 min, and (B3) 6.77 min match the theoretical accurate mass and isotopic pattern of a tetrachlorinated compound. The LC-Orbitrap MS analysis was performed in the negative polarity mode.

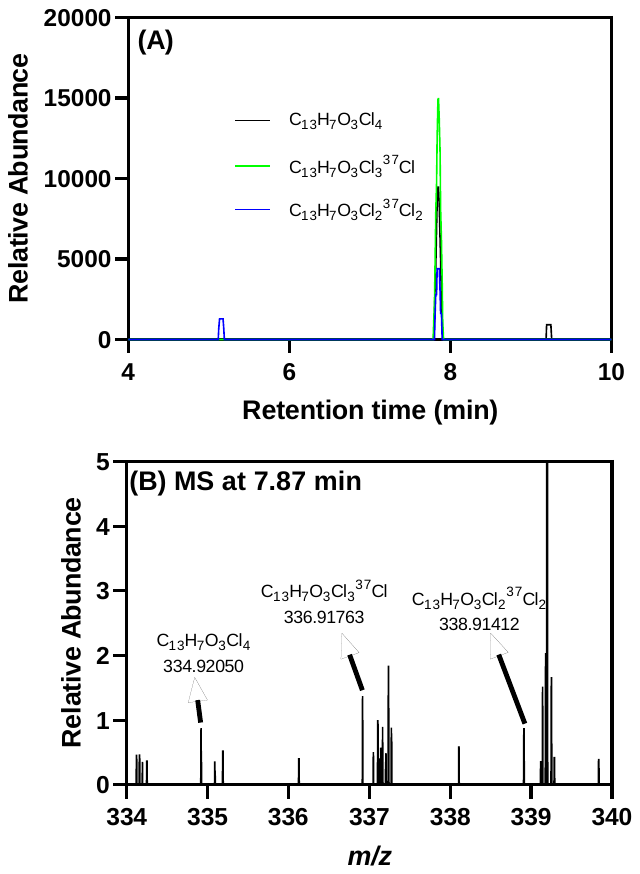

**Figure S7.** One tetrachlorinated MeO-OH-PCB was detected by LC-Orbitrap MS in the liver of PCB 52 exposed rats. (A) Chromatograms extracted based on the theoretical accurate mass of the top three high-abundance isotope ions of tetrachlorinated MeO-OH-PCBs ([C_13_H_7_O_3_Cl_4_]^-^, *m/z* 334.92056 for the monoisotopic ion) show a peak at 7.87 min. (B) The accurate masses of three high-abundance isotope ions at 7.87 min match the theoretical accurate mass and isotopic pattern of a tetrachlorinated compound. The LC-Orbitrap MS analysis was performed in the negative polarity mode.

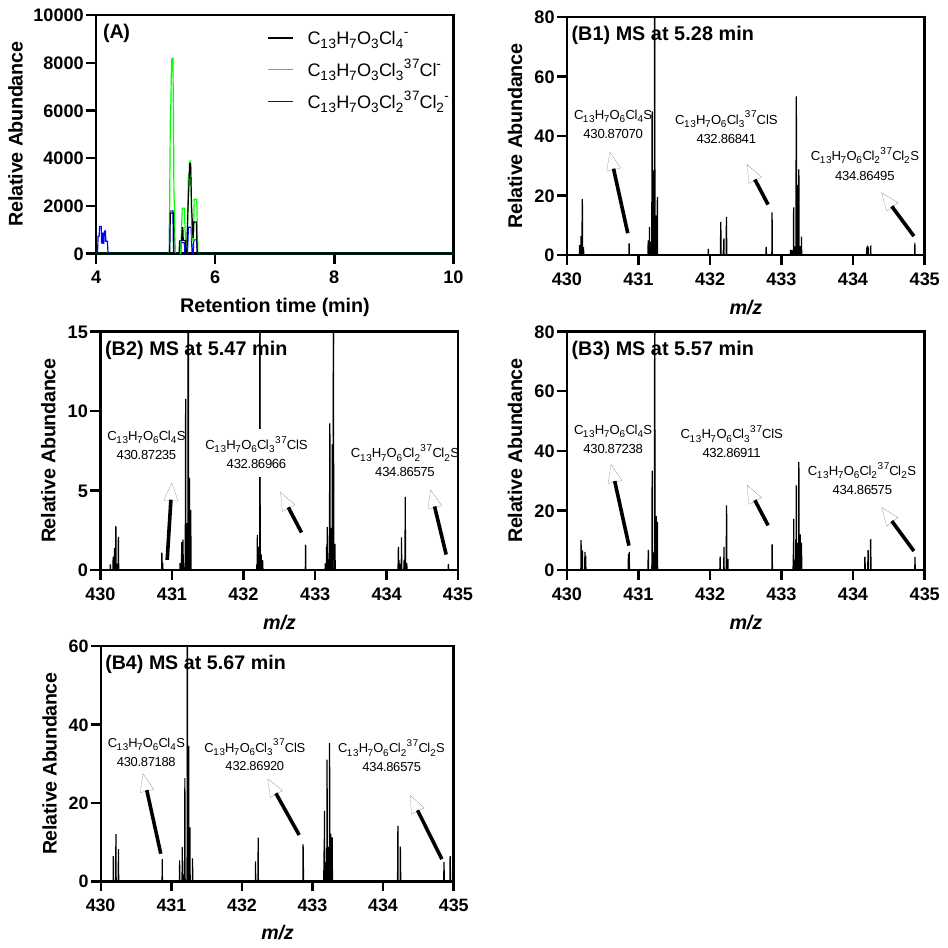

**Figure S8.** Four tetrachlorinated MeO-OH-PCB sulfates were detected by LC-Orbitrap MS in the liver of PCB 52 exposed rats. (A) Chromatograms extracted based on the theoretical accurate mass of the top three high-abundance isotope ions of a tetrachlorinated MeO-OH-PCB sulfate ([C_13_H_7_O_6_Cl_4_S]^-^, *m/z* 430.87229 for the monoisotopic ion) show peaks at 5.57, 5.47, 5.57, and 5.67 min. The accurate masses of four high-abundance isotope ions at (B1) 5.57 min, (B2) 5.47 min, (B3) 5.57 min, and (B4) 5.67 min match the theoretical accurate mass and isotopic pattern of a tetrachlorinated compound. The LC-Orbitrap MS analysis was performed in the negative polarity mode.

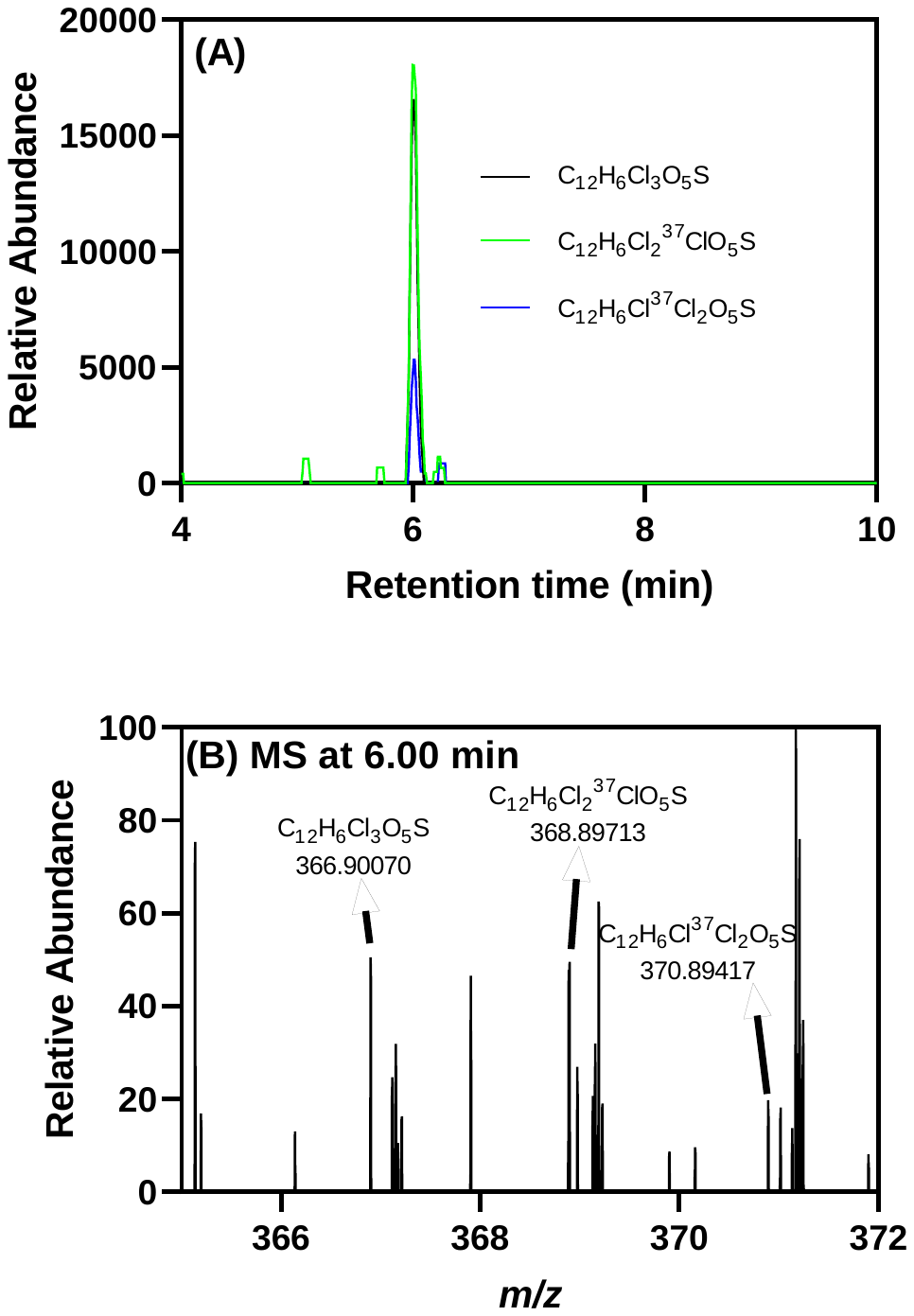

**Figure S9.** One trichlorinated OH-PCB sulfate was detected by LC-Orbitrap MS in the liver of PCB 52 exposed rats. (A) Chromatograms extracted based on the theoretical accurate mass of the top three high-abundance isotope ions of a trichlorinated OH-PCB sulfate ([C_12_H_6_C_l3_O_5_S]^-^, *m/z* 366.90070 for the monoisotopic ion) show a peak at 6.00 min. (B) The accurate masses of three high-abundance isotope ions at 6.00 min match the theoretical accurate mass and isotopic pattern of a trichlorinated compound. The LC-Orbitrap MS analysis was performed in the negative polarity mode.

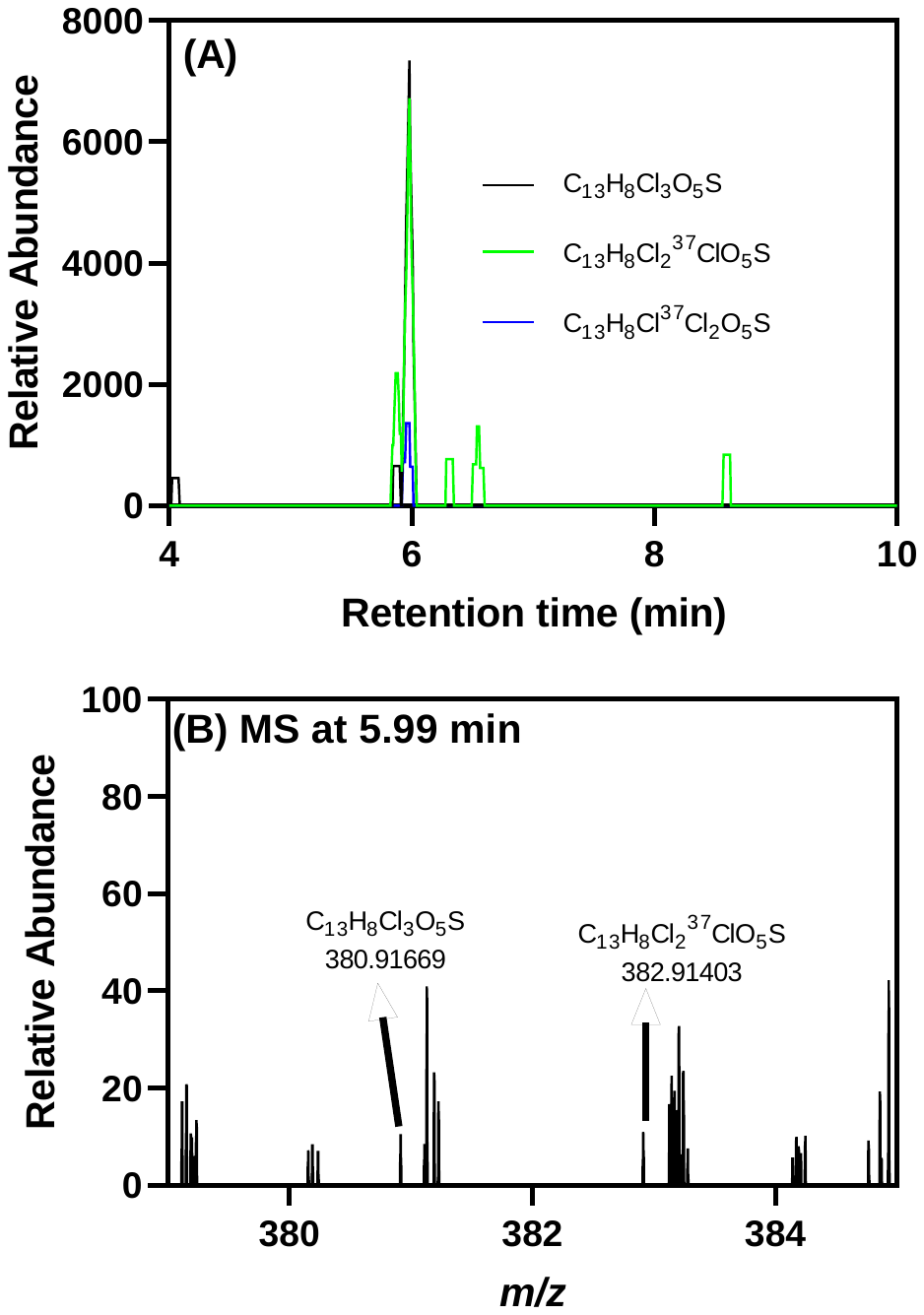

**Figure S10.** One trichlorinated MeO-PCB sulfate was detected by LC-Orbitrap MS in the liver of PCB 52 exposed rats. (A) Chromatograms extracted based on the theoretical accurate mass of the top two high-abundance isotope ions of a trichlorinated MeO-PCB sulfate ([C_13_H_8_Cl_3_O_5_S]^-^, *m/z* 380.91635 for the monoisotopic ion) show a peak at 5.99 min. (B) The accurate masses of three high-abundance isotope ions at 5.99 min match the theoretical accurate mass and isotopic pattern of a trichlorinated compound. The LC-Orbitrap MS analysis was performed in the negative polarity mode.

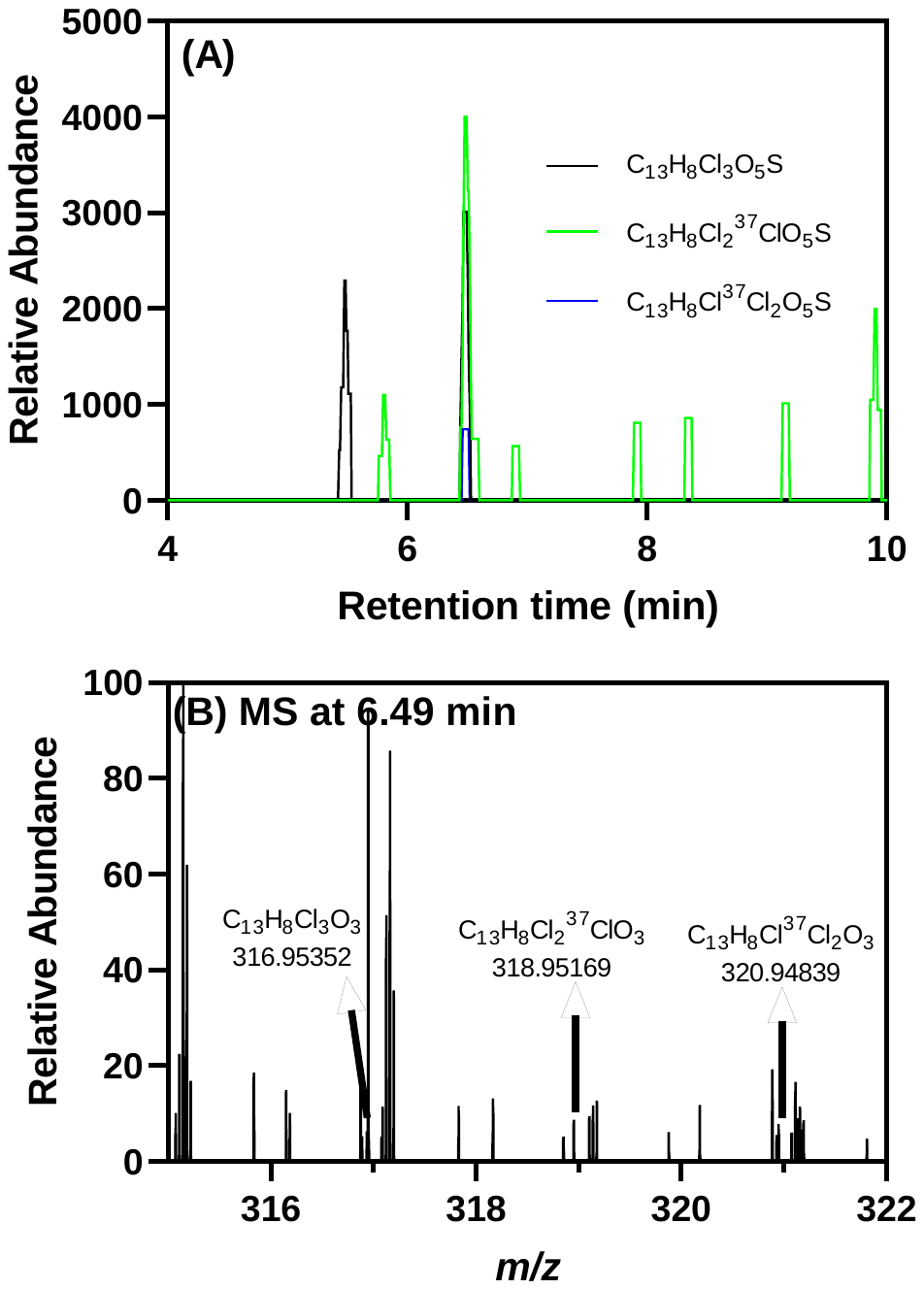

**Figure S11.** One trichlorinated MeO-diOH-PCB was tentatively detected by LC-Orbitrap MS in the liver of PCB 52 exposed rats. (A) Chromatograms extracted based on the theoretical accurate mass of the top three high-abundance isotope ions of a trichlorinated MeO-diOH-PCB ([C_13_H_8_Cl_3_O_3_]^-^, *m/z* 316.95445 for the monoisotopic ion) show a peak at 6.49 min. (B) The accurate masses of three high-abundance isotope ions at 6.49 min match the theoretical accurate mass and isotopic pattern of a trichlorinated compound. The LC-Orbitrap MS analysis was performed in the negative polarity mode.

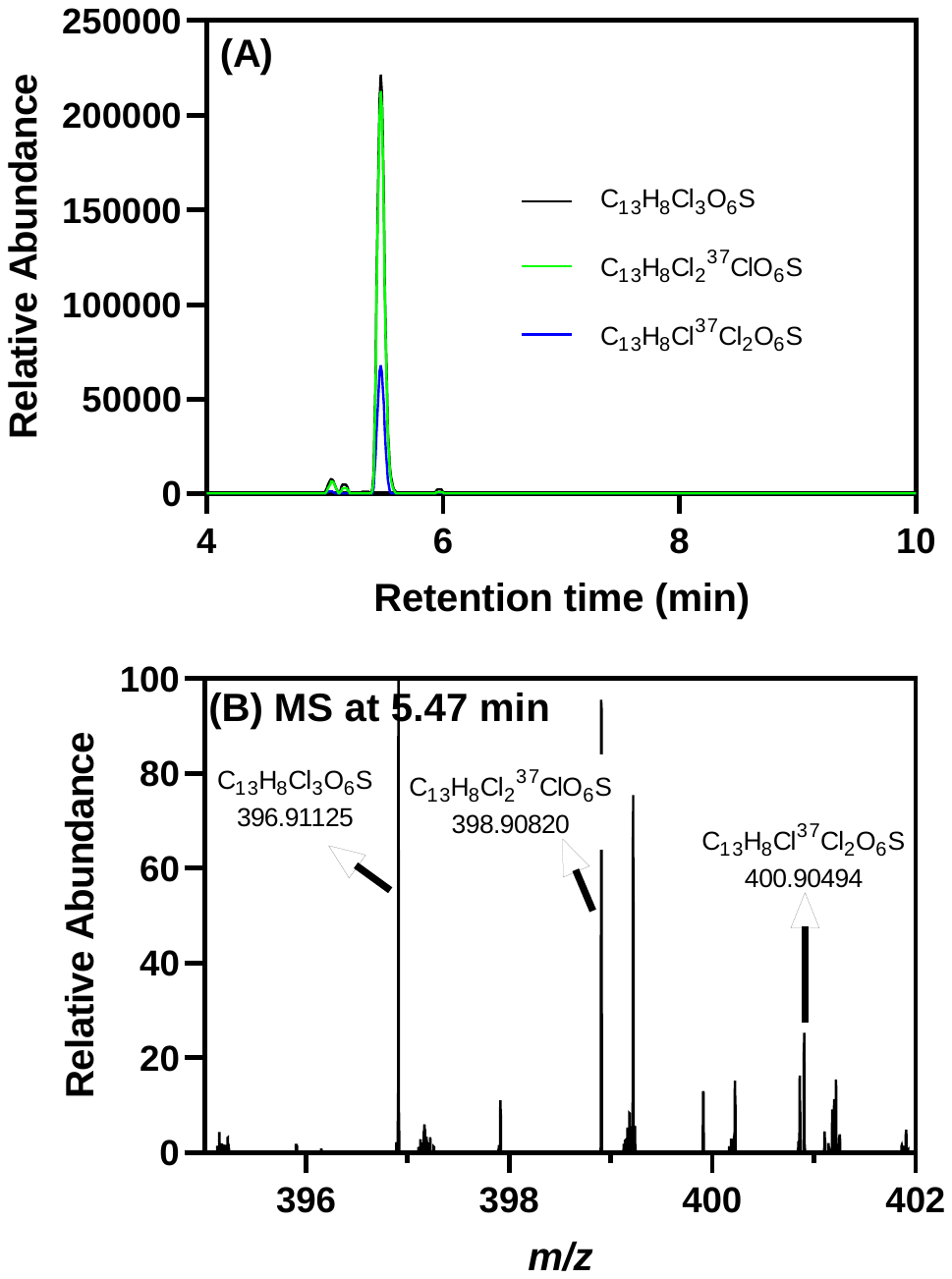

**Figure S12.** One trichlorinated MeO-OH-PCB sulfate was detected by LC-Orbitrap MS in the liver of PCB 52 exposed rats. (A) Chromatograms extracted based on the theoretical accurate mass of the top three high-abundance isotope ions of a trichlorinated MeO-OH-PCB sulfate ([C_13_H_8_Cl_3_O_6_S]^-^, *m/z* 396.91127 for the monoisotopic ion) show a peak at 5.47 min. (B) The accurate masses of three high-abundance isotope ions at 5.47 min match the theoretical accurate mass and isotopic pattern of a trichlorinated compound. The LC-Orbitrap MS analysis was performed in the negative polarity mode.

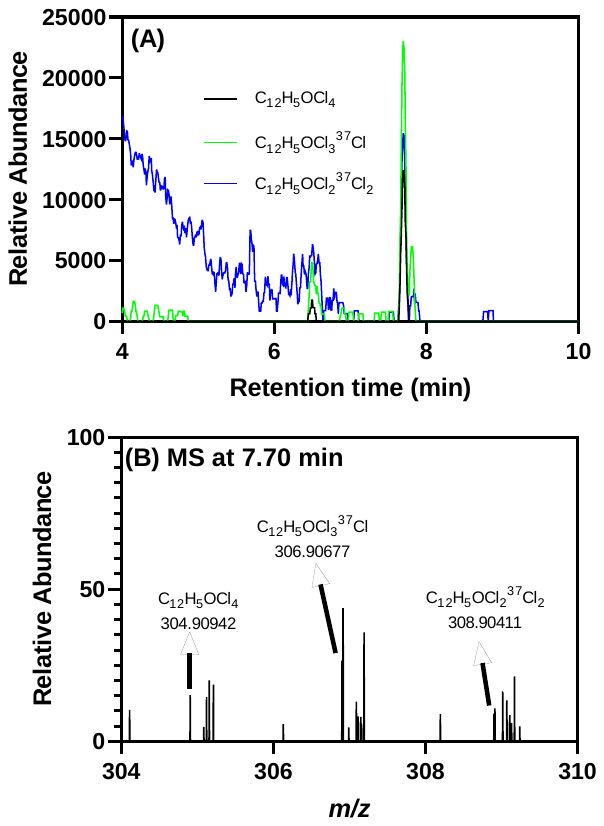

**Figure S13.** One tetrachlorinated OH-PCB was detected by LC-Orbitrap MS in the serum of PCB 52 exposed rats. (A) Chromatograms extracted based on the theoretical accurate mass of the top three high-abundance isotope ions of a tetrachlorinated OH-PCB ([C_12_H_5_OC_l4_]^-^, *m/z* 304.91000 for the monoisotopic ion) show a peak at 7.70 min. (B) The accurate masses of three ions at 7.70 min match the theoretical accurate mass and isotopic pattern of a tetrachlorinated compound. The The LC-Orbitrap MS analysis was performed in the negative polarity mode.

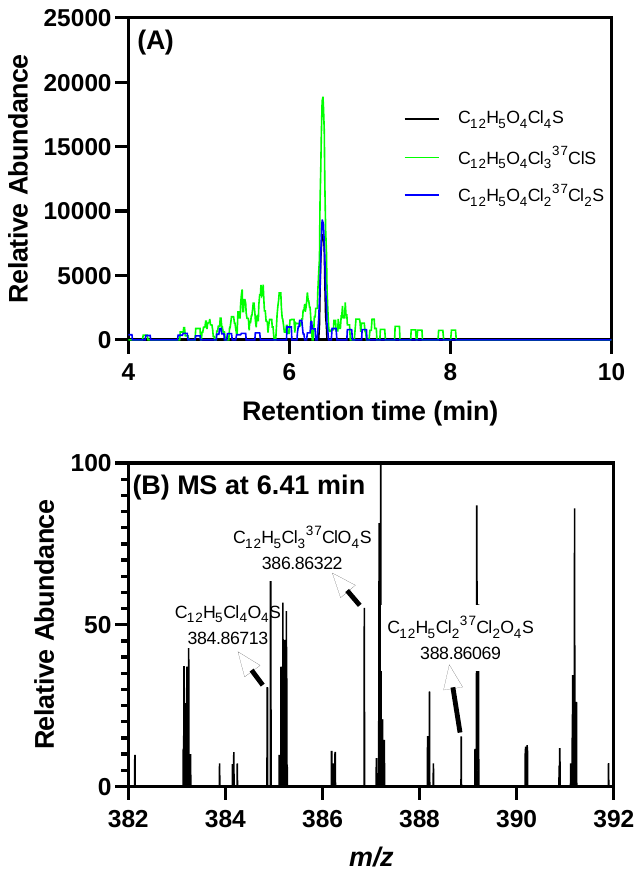

**Figure S14.** One tetrachlorinated PCB sulfate was detected by LC-Orbitrap MS in the serum of PCB 52 exposed rats. (A) Chromatograms extracted based on the theoretical accurate mass of the top three high-abundance isotope ions of a tetrachlorinated PCB sulfate ([C_12_H_5_Cl_4_O_4_S]^-^, *m/z* 384.86681 for the monoisotopic ion) show a peak at 6.41 min. (B) The accurate masses of three high-abundance isotope ions at 6.49 min match the theoretical accurate mass and isotopic pattern of a tetrachlorinated compound. The LC-Orbitrap MS analysis was performed in the negative polarity mode.

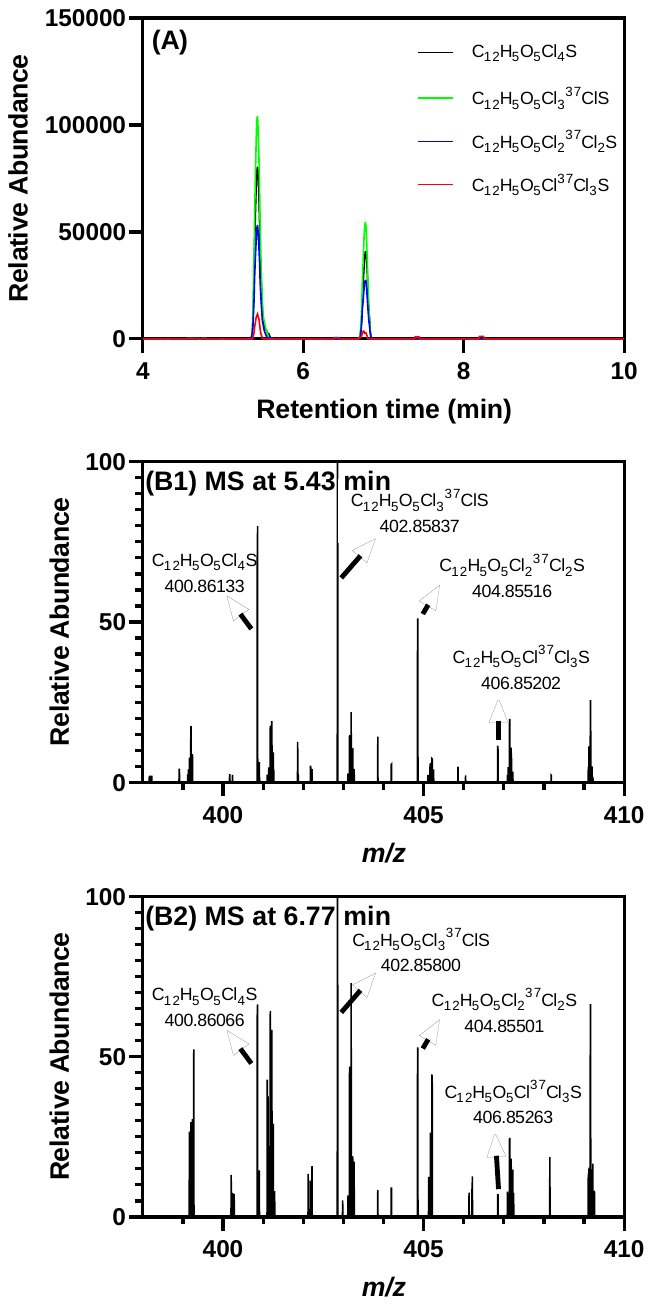

**Figure S15.** Two tetrachlorinated OH-PCB sulfates were detected by LC-Orbitrap MS in the serum of PCB 52 exposed rats. (A) Chromatograms extracted based on the theoretical accurate mass of the top four high-abundance isotope ions of a tetrachlorinated OH-PCB sulfate ([C_12_H_5_O_5_Cl_4_S]^-^, *m/z* 400.86173 for the monoisotopic ion) show peaks at 5.43 and 6.77 min. The accurate masses of four high-abundance isotope ions at (B1) 5.43 min and (B2) 6.77 min match the theoretical accurate mass and isotopic pattern of a tetrachlorinated compound. The LC-Orbitrap MS analysis was performed in the negative polarity mode, as described in the Supporting Information.

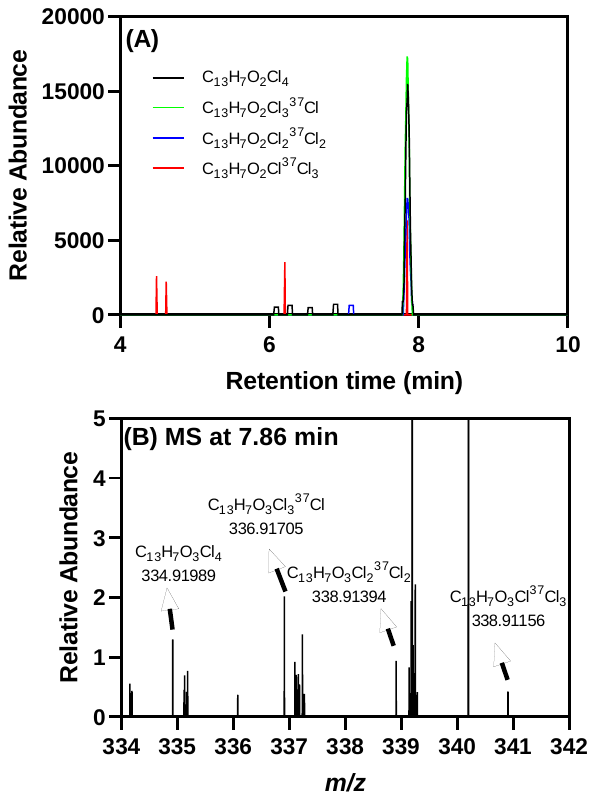

**Figure S16.** One tetrachlorinated MeO-OH-PCB was detected by LC-Orbitrap MS in the serum of PCB 52 exposed rats. (A) Chromatograms extracted based on the theoretical accurate mass of the top three high-abundance isotope ions of a tetrachlorinated MeO-OH-PCB ([C_13_H_7_O_2_Cl_4_]^-^, *m/z* 334.92056 for the monoisotopic ion) show a peak at 7.86 min. (B) The accurate masses of four high-abundance isotope ions at 7.86 min match the theoretical accurate mass and isotopic pattern of a tetrachlorinated compound. The LC-Orbitrap MS analysis was performed in the negative polarity mode.

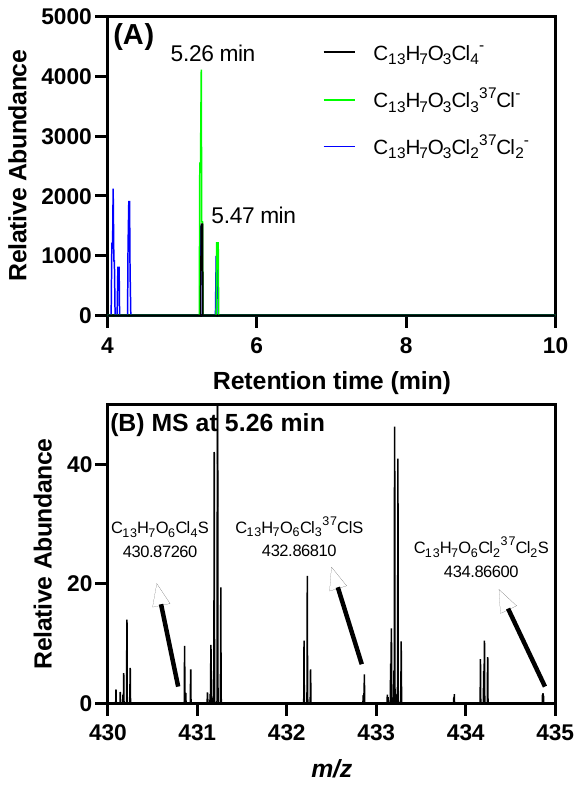

**Figure S17.** One tetrachlorinated MeO-OH-PCB sulfate was tentatively detected by LC-Orbitrap MS in the serum of PCB 52 exposed rats. (A) Chromatograms extracted based on the theoretical accurate mass of the top three high-abundance isotope ions of two tetrachlorinated MeO-OH-PCB sulfates ([C_13_H_7_O_6_Cl_4_S]^-^, *m/z* 430.87260 for the monoisotopic ion) indicate peaks at 5.26 min. (B) The accurate masses of several high-abundance isotope ions at 5.26 min match the theoretical accurate mass and isotopic pattern of a tetrachlorinated compound. The LC-Orbitrap MS analysis was performed in the negative polarity mode.

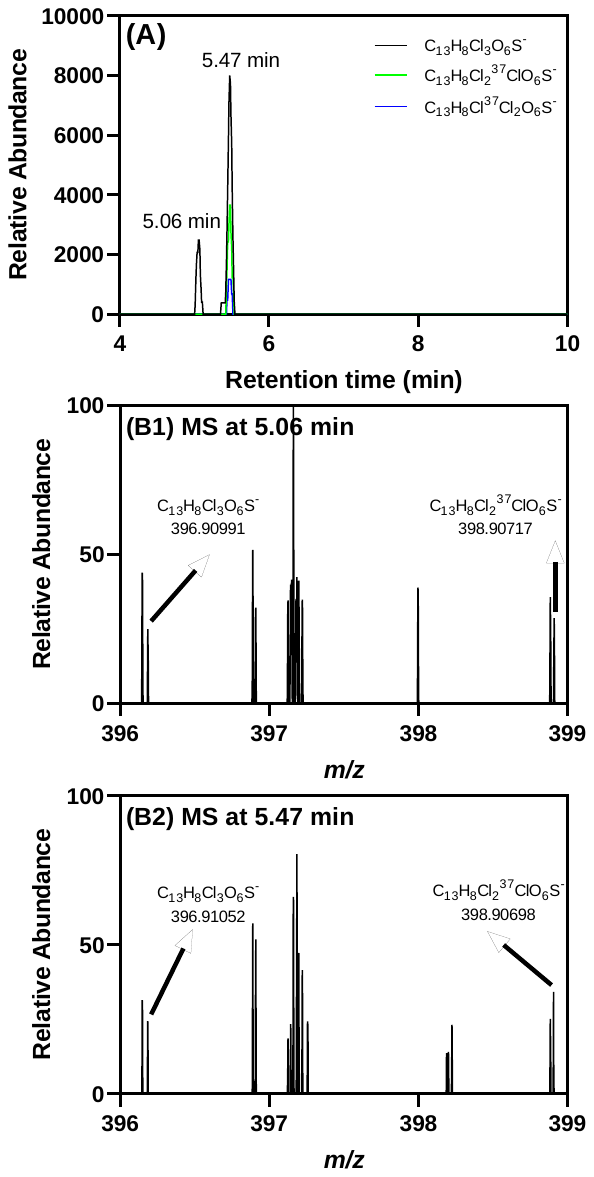

**Figure S18.** Two trichlorinated MeO-OH-PCB sulfates were tentatively detected by LC-Orbitrap MS in the serum of PCB 52 exposed rats. (A) Chromatograms extracted based on the theoretical accurate mass of the top two high-abundance isotope ions of trichlorinated MeO-OH-PCB sulfates ([C_13_H_8_C_l3_O_6_S]^-^, *m/z* 396.90991 for the monoisotopic ion) show peaks at 5.06 min and 5.47 min. The accurate masses of two high-abundance isotope ions at (B1) 5.06 min and (B2) 5.47 min match the theoretical accurate mass and isotopic pattern of a trichlorinated compound; however, the relative abundance of other isotopic peaks is too low to be shown. The LC-Orbitrap MS analysis was performed in the negative polarity mode.
